# The genetic variation landscape of African swine fever virus reveals frequent positive selection and adaptive flexibility

**DOI:** 10.1101/2020.08.12.249045

**Authors:** Yun-Juan Bao, Junhui Qiu, Yuzi Luo, Fernando Rodríguez, Hua-Ji Qiu

## Abstract

African swine fever virus (ASFV) is a lethal disease agent that causes high mortality in swine population and devastating loss in swine industries. The development of efficacious vaccines has been hindered by the gap in knowledge concerning genetic variation of ASFV and the genetic factors involved in host adaptation and virus-host interactions. In this study, we performed a meta-genetic study of ASFV aiming to profile the variation landscape and identify genetic factors with signatures of positive selection and relevance to host adaptation. Our data reveals a high level of genetic variability of ASFV shaped by both diversifying selection and selective sweep. The selection signatures are widely distributed across the genome with the diversifying selection falling within 29 genes and selection sweep within 25 genes, highlighting strong signals of adaptive evolution of ASFV. Further examination of the sequence properties reveals the link of the selection signatures with virus-host interactions and adaptive flexibility. Specifically, we discovered a site at 157th of the key antigen protein EP402R under diversifying selection, which is located in the cytotoxic T-cell epitope related with the low level of cross-reaction in T-cell response. Importantly, two multigene families MGF360 and MGF505, the host range factors of ASFV, exhibit divergent selection among the paralogous members, conferring sequence pools for genetic diversification and adaptive capability. By integrating the genes with selection signatures into a unified framework of interactions between ASFV and hosts, we showed that the genes are involved in multiple processes of host immune interaction and virus life cycles, and may play crucial roles in circumventing host defense systems and enhancing adaptive fitness. Our findings will allow enhanced understanding of genetic basis of rapid spreading and adaptation of ASFV among the hosts.

## Introduction

African swine fever virus (ASFV) is the causative agent of haemorrhagic fever in swine. ASFV mainly replicates in swine macrophages, causing up to 100% mortality rates in domestic pigs. However, the transmission pathway of ASFV is highly intertwined through the sylvatic cycle and domestic cycle involving multiple intermediate points, such as warthogs, soft ticks, wild boars, domestic pigs and human activities (Sánchez-Cordón et al., 2018). ASFV is thought to originate from and circulate in wild swine and soft ticks in Eastern Africa, and the first infection in domestic pigs was reported in Kenya in 1921 (Montgomery, 1921). From Africa, ASFV has spread to Europe in 1957 and 1960 (Portugal) and to Georgia in 2007 (Rebecca et al., 2008). This last introduction led to the expansion of the disease through the Caucasus and Russia (Oganesyan et al., 2013) to European Union countries, such as Estonia, Latvia, and Poland (Gallardo et al., 2014; Stokstad, 2017). More recently, ASFV was detected in wild boars in Belgium and an outbreak in a pig farm was reported in China in mid-2018 (Garigliany et al., 2019; Ge et al., 2018). Since then, the virus has continuously spread to neighboring countries in Asia, affecting hundreds of millions of swine population (FAO, 2019).

Since there is no commercially available vaccine against ASFV infection, current disease control is based on physical quarantine and animal slaughtering. A large amount of them have been killed since the spread of infection globally, causing substantial damages on the swine population. Development of efficacious therapeutic and prophylactic tools has been largely hindered by the limited knowledge of genetics properties and the evolutionary adaptation of this highly pathogenic virus.

ASFV is a large double-stranded DNA (dsDNA) virus with a genome length of 170∼194 kb. Tens of genomes of ASFV strains have been completed by using high-throughput sequencing technologies. Previous studies mainly focused on a limited number of virulence determinants or host-range factors, such as EP402R (Borca et al., 1998), EP153R (Hurtado et al., 2011), A238L (Powell et al., 1996), and the highly variable multigene families (MGFs) at both ends of the genome (Chapman et al., 2008; de Villiers et al., 2010; Dixon et al., 2013). Specifically, studies using engineered deletion mutants investigated the variation patterns of MGF genes (Chapman et al., 2008; Rodríguez et al., 2015), showing that MGF genes are relevant to host interactions and might be responsible for host range functions (Dixon et al., 2013; Zsak et al., 2001).

However, there are a limited number of studies on systematic characterization of genetic properties for the whole gene set in the genome-wide scale (Chapman et al., 2008; de Villiers et al., 2010). As a dsDNA virus, ASFV has an estimated substitution rate μ ∼ 6.7x10^-4^ (substitutions per site per year) (Michaud et al., 2013), roughly between that of RNA viruses such as the influenza virus with μ ∼ 10^-3^ (Hanada et al., 2004) and that of other large dsDNA viruses such as herpes simplex type I virus with μ ∼ 10^-5^ (Duffy et al., 2008). This substitution level is much higher than that of many bacterial species such as *Streptococcus pneumoniae* with μ ∼ 10^-6^ (Croucher et al., 2013). The high substitution rate indicates a high level of variability in the seemingly conserved central regions previously thought. This high level of genetic variability may have important implication for understanding the puzzling adaptive capability and host range of ASFV. For instance, investigation of the breath of the variability on the genome-wide scale will help to understand the molecular basis underlying the evolutionary adaptation; identifying the selection pressures acting on the genes will reveal the genetic factors exposed to host-virus interfaces; dissecting the genetic diversity of MGF genes will help to clarify the driving force of sequence divergence and functional diversification of MGF. During the decades of study on ASFV, those critical issues are still largely unknown.

In this study, we performed a meta-genetic study by using sophisticated computational methods to profile the variation landscape of ASFV and identify the genetic factors under positive selection aiming to characterize the genetic factors relevant to the versatile adaptation and host interactions for ASFV.

## Materials and Methods

### Comparative genomic study and phylogenetic inference

The genomic sequences and annotations of ASFV used in this study were downloaded from NCBI GenBank (ftp://ftp.ncbi.nlm.nih.gov). The non-redundant genomes were identified and used for downstream analysis by excluding those with close evolutionary distance (< 0.001 substitutions per site), the same isolation countries and isolation time with other strains (see Table S1). The core genome represents the genes or genomic locations present in all studied strains of a species and we created the core genome of ASFV by aligning the shredded genomes against the reference strain Georgia-2007 and extracting the genomic regions mapped by all other genomes. Finally, the core genome contains 139,677 base pairs and was used for single nucleotide polymorphism (SNP) detection. The bases at the variant loci for each ASFV genome were concatenated for distance estimation and phylogeny construction using MEGA6 (Tamura et al., 2013) and SplitsTree (Huson and Bryant, 2006). The pair-wise distance was measured by substitutions per site with the model of maximum composite likelihood and the tree topology was inferred using the Neighboring-Joining method with a bootstrap value of 1,000. The tree was also constructed using the Maximum Likelihood method. The tree topologies are consistent between different methods. Tajima’s *D* is a statistic for testing the neutrality of the mutations on the overall scale by computing the difference between the average number of pairwise nucleotide differences and the number of segregating sites and was calculated as defined by Tajima (Tajima, 1989).

### Detection of functional domains

The functional domains of the genes were detected by comparison with the PFAM database (Punta et al., 2012). The hits with score ≥ 20 or *E*-value ≤ 0.003 were considered to be significant and tabulated.

### Generation of pan-genome and orthologous groups of ASFV

The pan-genome of a species is the whole set of genes encoded by all studied strains. The genes present in different strains facilitating similar functions form orthologous groups and are key components of the pan-genome. On the other hand, paralogous genes are those duplicated in the same strain from a common ancestor with related but divergent functions (Jensen, 2001). Derivation of a new paralogous member in a gene family will lead to emergence of new functions and expansion of the pan-genome size. The pan-genome of the 27 non-redundant ASFV genomes was generated using Roary yielding 192 pan-genes encoded by at least one strain of ASFV (Page et al., 2015). The amino acid translation of the pan-genes were aligned against each ASFV genome using BLAST tblastn in order to determine the 5’- and 3’-end of the pan-genes in each genome and rescue the genes interrupted by point mutations. Only the genes present in more than 70% of the 27 non-redundant genomes were cataloged into orthologous groups and considered for downstream positive selection detection. The orthologous groups of MGF genes were refined by stratifying the tandem locations of the paralogous members in each genome to avoid mis-classification given the fact some MGF genes have higher similarities with paralogs than orthologs. The fusion genes were not considered for further analysis.

### Analysis of selection pressures on the ASFV genes

Multiple sequence alignment was performed at first in amino acids for each orthologous gene group and then were back converted to alignment in nucleotides. All the alignments were manually curated to make the coding sequences in frame. The calculation of non-synonymous substitutions dN and synonymous substitutions dS was based on the Nei & Gorojobri model (Nei and Gojobori, 1986). Likelihood ratio tests (LRT) of selection pressures acting on individual sites of ASFV genes were carried out using PAML with the site-specific model (Yang, 2007). For each gene, two LRT tests were conducted, *i.e.*, M2 versus M1 and M8 versus M7. The genes with *p*-value ≤ 0.05 for the test between M8 versus M7 were considered to contain signals with significant positive selection. Only the sites showing positive selection with a posterior probability ≥ 0.9 in M8 were tabulated. The posterior probability was calculated using PAML with the Bayes empirical tests (Yang et al., 2005). Likelihood ratio tests of divergent selection of MGF genes were performed using the branch-site Model A in PAML (Zhang et al., 2005). A total of 13 pairs of paralogous members from MGF360 (1L:2L, 1L:3L, 2L:3L, 4L:6L, 8L:10L, 8L:13L, 10L:13L, 9L:11L, 9L:12L, 11L:12L, 14L:16R, the ancestral branch of 1L/2L:3L, and the ancestral branch of 4L/6L:16R) and 13 pairs from MGF505 (1R:4R, 1R:5R, 4R:5R, 2R:4R, 2R:5R, 1R:2R, 2R:10R, 9R:10R, 6R:7R, 6R:9R, 7R:9R, 6R:10R, and 7R:10R) were chosen for LRT of Model A. Either member in the pairs was treated as foreground for the Model A test. The sites under positive selection with a posterior probability ≥ 0.8 for MGF360 and ≥ 0.9 for MGF505 using Bayes empirical tests were tabulated and mapped to the respective secondary structures.

### Multiple sequence alignments of orthologs and paralogs of the MGF genes

Since sequence similarities between orthologs of MGF genes are much higher than that of paralogs (except MGF360-1L and 2L, MGF505-6R and 7R), we performed multiple sequence alignment in amino acids at first for orthologous members of each paralog of MGF and then for paralogous groups of all MGF360 (except 15R, 18R, 19R, 21R and 22R), or MGF505 (except 3R and 11L due to the high divergence with other paralogs and low reliability of alignment). The alignments in amino acids were back converted to multiple alignments in nucleotides.

### Secondary structure prediction

The secondary structures of B475L and MGF300-4L were predicted using PSIpred (Buchan and Jones, 2019), and those of MGF360 and MGF505 using PROMALS3D (Pei et al., 2008).

### Tertiary structure prediction and structure-guided sequence alignment

The tertiary structure of EP402R was modeled using PHYRE server with the structure of human CD2 as template (Kelley et al., 2015). Multiple sequence alignment of EP402R and its homologs in animals, including human CD2 (Bodian et al., 1994) (PDB ID: 1hnf), human CD58 (Ikemizu et al., 1999) (PDB ID: 1ccz), rat CD2 (Jones et al., 1992) (PDB ID: 1hng), rat CD48 (Evans et al., 2006) (PDB ID: 2dru), and boar CD2 (modeled with PHYRE server) was guided by the tertiary structures. The graphical presentation of the alignment was prepared using Espript (Robert and Gouet, 2014). The structures of the proteins were presented and analyzed using PyMOL (Benoit et al., 2008).

### Statistical analysis

The statistical tests used in this study including Hypergeometric test, Mann-Whitney U-test, T-test, and Chi-squared test were performed in the R environment.

### Identification of regions with selective sweep

The population size is highly unbalanced between the two subpopulations α (21 strains) and β (5 strains), therefore we at first identified the SNPs associated with between-population subdivision and within-population homogeneity for the clade α and β by selecting loci with the major allele frequency > 85% in clade α and alternative allele frequency > 80% in clade β. The selected SNPs were subject to detection of selective sweep using the clustering algorithm described in (Bao et al., 2016). Briefly, a non-synonymous SNP is randomly chosen in a specific gene as the initial cluster and each initial cluster is then iteratively extended until its spanning range approaches the specified sweep length or the boundary of the gene or gene operon. The cluster is further extended to merge the neighboring SNPs or clusters by minimizing the root-mean-square of inter-SNP distances. The significance of the clustering for each cluster with *m* distinct SNPs spanning a length of L was evaluated using the gamma distribution with the mean SNP rate μ as the rate parameter under the null hypothesis that the SNPs are randomly and independently distributed on the genome:

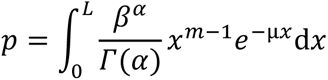

## Results

### Single nucleotide polymorphism (SNP) detection and selection pressure in the core genome of ASFV

We performed comparative genomic study of the ASFV strains by aligning the genomic sequences of the strains to the core genome. The list of ASFV genomes we used is shown in Table S1. Using 27 non-redundant genomes, we identified 18,070 SNPs, of which 6088 are non-synonymous, corresponding to an average of 129 SNPs/kb. In order to examine the influence on variation detection from the five distantly evolved strains from Africa, *i.e.*, Ken05-Tk1, Kenya-1950, Ken06-Bus, UgandaN10-2015, and UgandaR7-2015 (Fig. 1), we excluded the five strains, repeated the comparative analysis and obtained 12,652 SNPs with an average 91 SNPs/kb, again reflecting the high genetic diversity of ASFV. The high mutation rate is in contrast with the previous notion of high conservation of the core genomes of ASFV. Therefore, we further estimate the overall selection pressure exerted on the ASFV population using Tajima’s *D* test (Tajima, 1989). The calculation of Watterson’s estimator *θ* (Watterson, 1975) gives a genome-wide average mutation rate of 0.025, significantly greater than the average pair-wise nucleotide difference of 0.019. It results in a negative Tajima’s *D* value of -2.30, indicating evolutionary positive selection of the ASFV population.

**Fig. 1.**
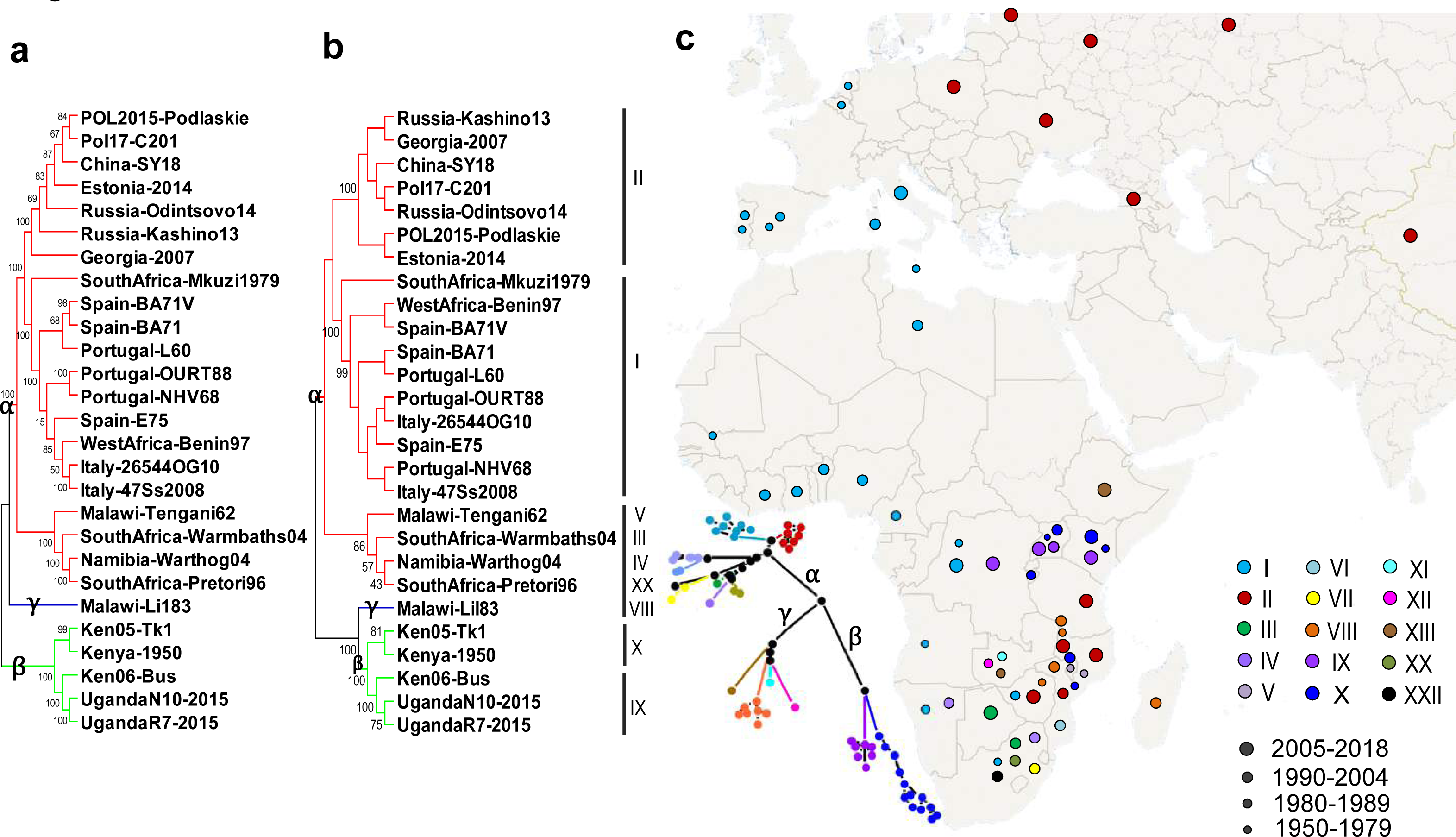
Phylogenetic tree and geographical distribution of ASFV strains. (a) Phylogeny built from the core genome of 27 non-redundant ASFV strains. (b) Phylogeny built from the full-length structural gene p72 (B646L) of the 27 non-redundant ASFV genomes. The subtypes are shown on the right. (c) Geographical distribution of 85 non-redundant ASFV isolates and the phylogeny constructed using the C-terminal 414 bp of p72 sequences available from public databases. The partial p72 sequences of the 85 non-redundant ASFV isolates with unique geographical location and isolate time were compiled from the NCBI database https://www.ncbi.nlm.nih.gov/ and mapped to the geographical locations. The trees were inferred using the Neighboring-Joining method with 1000 bootstrap. The trees built from all three datasets forms three major clades α, β, and γ indicated on the corresponding branches.

### Phylogenetic structure of the ASFV population

The genome-wide phylogeny was inferred using the core genome SNPs of the 27 non-redundant strains (Fig. 1a and Supplementary file 1a). The phylogenetic tree identifies three major distantly related clades (α, β, and γ). The three-clade topology is consistent with that derived from the full-length structural gene p72 (B646L) of the same set of genomes and the partial-length p72 sequences from a broader set of 85 isolates (Fig. 1b,c, Fig. S1, and Supplementary file 1b). The first clade α contains three closely related subgroups, comprising isolates from Europe of genotype I, isolates from Caucasus of genotype II, and isolates from Southern Africa of diverse genotypes, respectively. The second clade β consists of isolates from Eastern Africa of genotype X and IX, which are the predominant genotypes causing outbreaks in this area (Atuhaire et al., 2013). The third clade γ mainly contains Eastern African isolates of genotype VIII, XI, XII, and XIII, although only one complete genome is available in this clade (Malawi-Lil83 of genotype VIII). The phylogeny topology is consistent with that constructed previously based on different number of ASFV strains (de Villiers et al., 2010; Rebecca et al., 2008).

We observed two prominent features of the phylogenetic structure and geographical distribution depicted in Fig. 1. First, the tree has a total branch length of 2.1 substitutions per site. The long phylogenetic distance and relatively short separation time between the three clades, especially α and β indicates that they have accumulated a significant number of genetic differences in a short period of time. Secondly, the virus has recurrently emerged at the same countries at different time points but exhibits significant genomic modifications, such as those isolates from Malawi (Malawi-Tengani62 and Malawi-Lil83 with a genetic distance of 0.09 substitutions per site). It re-elaborates the rapid adaptation of ASFV to host environments and the complexity of the transmission pathways of ASFV. Third, no temporal-spatial dynamics pattern can be inferred from the phylogenetic structure except the recent spreading of genotype II strains. Next, we will investigate in details the genetic variation profile of the whole population of ASFV, but without focusing on specific genotypes.

### Identification of genes with high frequencies of non-synonymous mutations

The pattern of gene duplication and loss affecting the MGFs at both ends of the ASFV genomes has been intensively studied (Donnell et al., 2015; Krug et al., 2015; Rodríguez et al., 2015), largely due to the postulated roles of MGF360 and MGF505 in host immune evasion and infection tropism (Dixon et al., 2013; Donnell et al., 2015). Here, we focus on the whole genome to characterize the genetic variation properties. We at first identify the variations associated with virulent phenotypes of ASFV strains. The low number of non-virulent strains in the currently known data set prevents us from performing a robust statistical association study, we quantified the non-synonymous allelic changes uniquely present in the two natural isolates with low virulence, *i.e.*, Portugal-NHV68 and Portugal-OURT88. A total of 13 non-synonymous mutations from 10 genes were uniquely present in the two Portugal isolates (Table S2). However, none of the genes is enriched with the unique mutations with statistical significance in comparison with the genome-wide average using Hypergeometric tests.

Therefore we further examined the distribution of all 6088 non-synonymous mutations along the genome and identified the gene loci mutated more frequently than the genome-wide average (Fig. S2a). The analysis using Hypergeometric test ranked 23 genes to be significantly enriched with non-synonymous mutations (multiple testing corrected *p*-value ≤ 0.001) but not with synonymous mutations (multiple testing corrected *p*-value ≥ 0.05) (Table S3 and Fig. S2b). Half of the genes are the members of MGF360, MGF505, and MGF300. The list also includes the genes involved in DNA replication/repair, nucleotide metabolism, redox pathway, host interactions, and others with unknown functions. The non-synonymous mutations in the 23 genes were further laid on each protein domain architecture identified by comparison with the PFAM database (Punta et al., 2012) (Table S4). We found no significant difference of the mutation distribution between the key functional domains and the neighboring regions. The zoomed-in view of the density distribution of the non-synonymous mutations along the domain architectures for the top genes is shown in Fig. 2c.

**Fig. 2.**
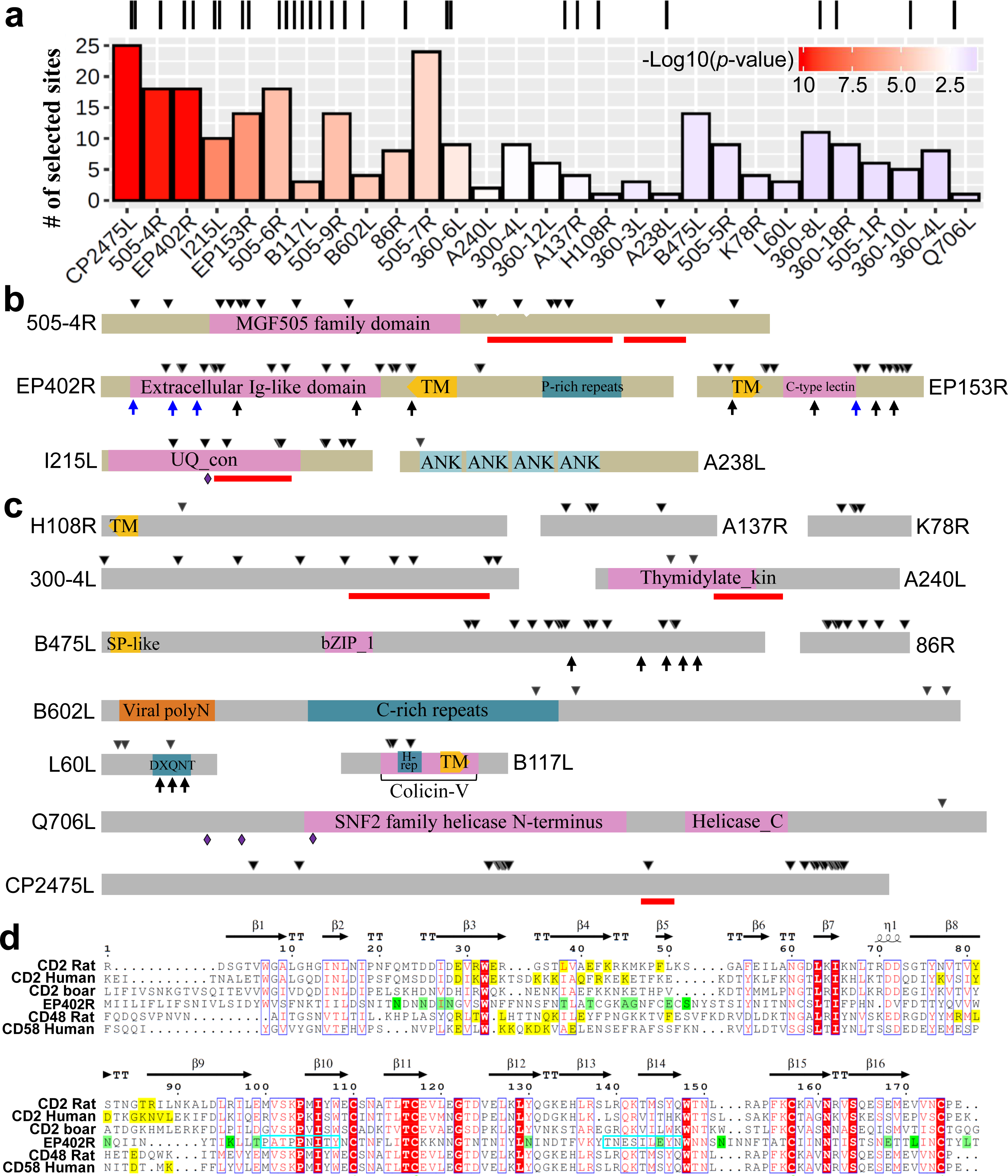
Genetic and functional properties of genes with positive diversifying selection signals. (a) The genes containing sites under positive diversifying selection (*p*-value ≤ 0.05). Top panel: the genomic locations of the genes. Bottom panel: histogram representation of the number of sites with significant selection in each gene (posterior probability ≥ 0.9). (b,c) Layout of the positively selected sites on the domain architectures of the key genes known to be relevant to host interactions (b) and of novel candidate genes with unknown host interactions (c). The positively selected sites (in black triangles) of EP402R, EP153R, MGF505-4R, B475L, and B117L are largely located in the variable regions or near around short repeat-rich regions (arrows, with blue ones for putative *N*-linked glycosylation sites). The functional domains are represented as colored bars and the transmembrane domains as directed frames pointing towards outside of the membrane. The active sites are shown as diamonds. The red bars show overlapping regions with signatures of selective sweep. The lengths of the proteins might be longer than the actual length due to gaps induced by multiple alignments. The length of the protein CP2475L is in a shrunk scale due to its exceptionally large size. Abbreviations: DXQNT: DXQNT repeats; TM: transmembrane domain; P-rich repeat: proline-rich repeats; ANK: ankyrin repeat; UQ_con: ubiquitin-conjugating enzyme; H-rep: histidine-rich repeats; Colicin-V: Colicin-V production domain; SP-like: signal peptide-like domain; Thymidylate_kin: thymidylate kinase domain; bZIP_1: basic leucine zipper domain; Viral polyN: viral polyprotein N-terminal domain. (d) Multiple sequence alignment of the extracellular Ig-like domain of EP402R and its homologs in rat (CD2, CD48), human (CD2, CD58), and boar (CD2). The secondary structure of rat CD2 is displayed on the top of the alignment with β strands in arrows and β turns in TT. The known ligand-binding sites of CD2, CD48, and CD58 are highlighted in yellow and the positively selected sites in EP402R are in green (posterior probability ≥ 0.9) or light green (posterior probability ≥ 0.8). Two known epitopes F3 and A6 in ASFV strain Spain-E75 are framed in cyan boxes.

### Identification of genes under positive selection based on the dN/dS method

The high rate of non-synonymous mutations observed prompted us to test the potential occurrence of positive diversifying selection acting on the ASFV-encoded genes. Positive diversifying selection is represented as elevated amino acid diversity within or across the populations resulting in selection of multiple phenotypes. It can be detected by measuring the rates of non-synonymous substitution (dN) and synonymous substitutions (dS) and calculating their ratio dN/dS. We at first calculated the dN/dS for each gene based on the Nei & Gojobori model (Nei and Gojobori, 1986). The analysis shows that most of the genes have a value of dN/dS < 0.5 and the average value of dN/dS is 0.1, revealing the evolutionary stability of the genes (Table S5). Notably, at the top of the list are six genes with the value of dN/dS ≥ 1 (D1133L, DP63R, 86R, EP153R, EP402R, and MGF505-4R). By removing three genes with deflated values of dS due to increased selection against synonymous substitutions (dS < 0.028, *p*-value < 0.02, one-tailed t-test), we finally obtained three genes (EP153R, EP402R, and MGF505-4R) with dN/dS > 1, subject to potential positive selection. Among them, the gene MGF505-4R with the value of dN/dS = 1.2 was also found to be significantly enriched with non-synonymous mutations in the previous section, implying strong positive selection acting on this gene. The other two genes, the CD2 homolog protein EP402R and C-type lectin-like protein EP153R, were previously shown to be involved in host immune evasion and the hemagglutination ability of ASFV depends on these two genes (Galindo et al., 2000; Ruiz-Gonzalvo et al., 1996).

### Test of selection pressures on individual sites of genes

In most organisms, the genes with dN/dS>1 are rare because non-synonymous mutations are generally detrimental to protein functions and are not preferred. Therefore, the individual sites positively selected are usually masked by the low average value of gene-wide dN/dS. In order to unravel the potential selection acting on specific sites of the genes, we performed likelihood ratio tests (LRTs) using the site-specific model of dN/dS (ɷ) in PAML (Yang, 2007). We identified 29 genes having been subject to potential positive diversifying selection (*p*-value ≤ 0.05, Chi-squared test) on an average of 3.1% (±2.4%) of sites (posterior probability ≥ 0.9) (Fig. 2a and Table S6). The list of genes under positive selection covers 11 of the 18 genes with *p*-value ≤ 0.05 and 8 of 10 genes with *p*-value ≤ 0.01 identified by a comparative study of 11 complete genomes (de Villiers et al., 2010).

The genes here we identified include 17 candidates known to be involved in host cell interactions, such as EP402R, EP153R and MGF genes. Notably, we also discovered twelve novel candidates, which have not been shown to be related with host interactions or investigated thoroughly experimentally, such as the highly divergent proteins B117L and B602L, and the conserved structural protein pp220/CP2475L (Table 1 and Table S6).

**Table 1.**
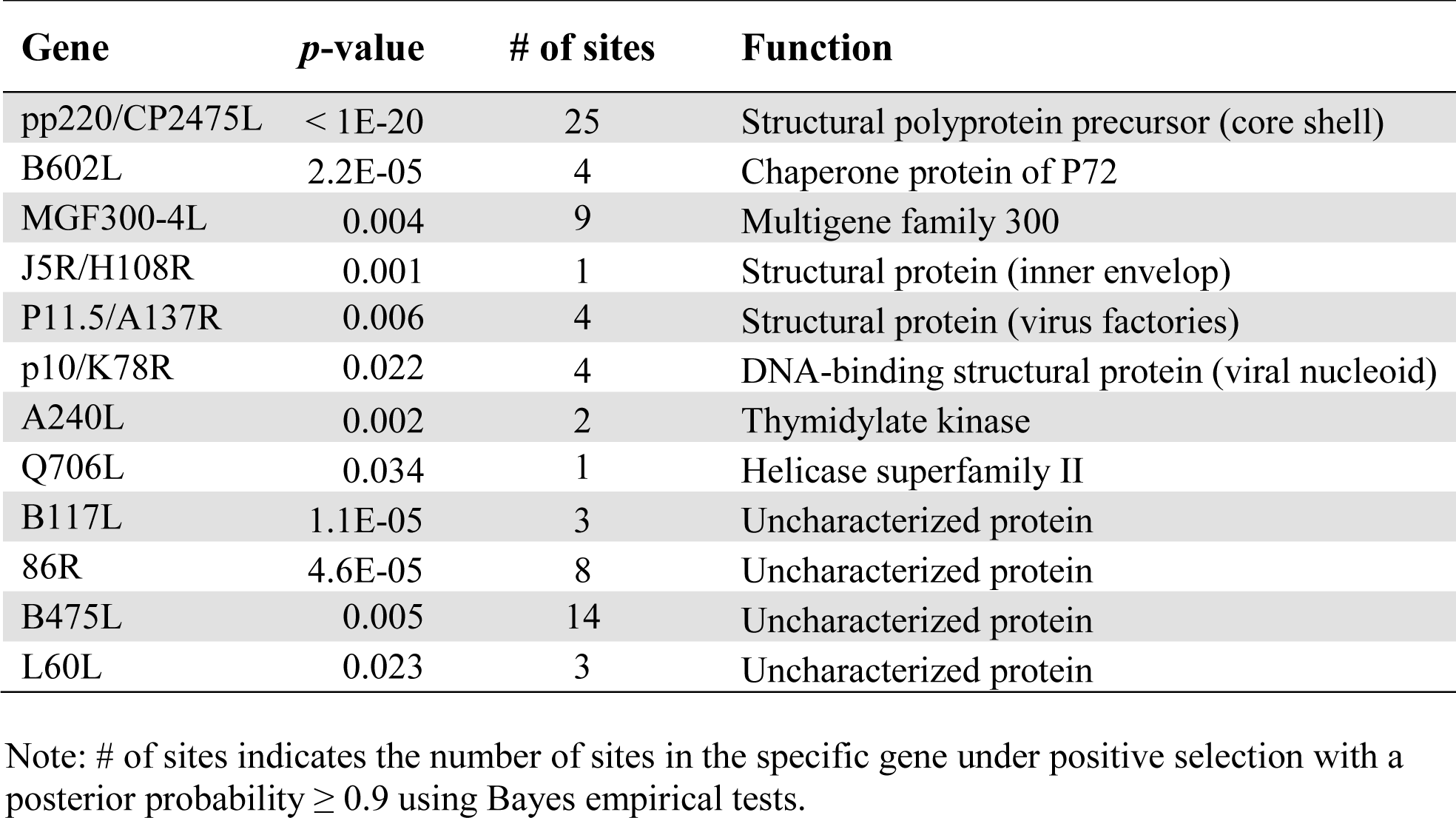
Novel candidates with positive selection signals at a fraction of sites with ⍵ (dN/dS) >1 based on the likelihood ratio tests.

### Functional implication of the positively selected sites

In order to ascertain the functional implication of the positively selected sites in the genes, we tabulated the sites under positive selection in each gene with a posterior probability ≥ 0.9 and mapped the sites to the domain architectures of the genes (Fig. 2b,c and Supplementary file 2). The positively selected sites are largely located in the variable regions or around the short repeats of the genes, such as EP402R, EP153R, B117L, and B475L. Specifically, eighteen positively selected sites are identified in EP402R and significantly enriched in the extracellular domain (*p*-value = 0.046, Hypergeometric test), which is highly variable among the ASFV lineages. The extracellular domain has an Ig-like structure resembling to host CD2 protein and is essential for binding of red blood cells to infected cells or extracellular virions (Alejo et al., 2018; Borca et al., 1998; Rodríguez et al., 1993). Here we use EP402R as an example to demonstrate the feasibility of using positively selected sites to delineate their links with virus-host interactions. We collected the CD2 homologs of EP402R in animals with known functions and structures, and performed structure-guided comparison with the EP402R extracellular Ig-like domain (Fig. 2d and Fig. 3). As a CD2 homolog, the extracellular domain of EP402R consists of a constant C-set and a variable V-set Ig-superfamily domain (Fig. 3a-d). We then mapped the positively selected sites to the aligned sequences and the tertiary structures. It is remarkable that the sites under positive selection predominantly reside in the loop regions on the top of the V-set domain of EP402R, in clear contrast with the location of the ligand-binding sites of host CD2 at the side face of the V-set domain (Fig. 3a-c) (Davis et al., 1998). The orthologous loop regions in Ig antibodies are responsible for facilitating specificity of antibodies to recognize antigens (Morea et al., 2000). It indicates the potential roles of the positively selected sites in the loop regions of EP402R in determining specificity of ASFV for host cell recognition and enhancing adaptability.

**Fig. 3.**
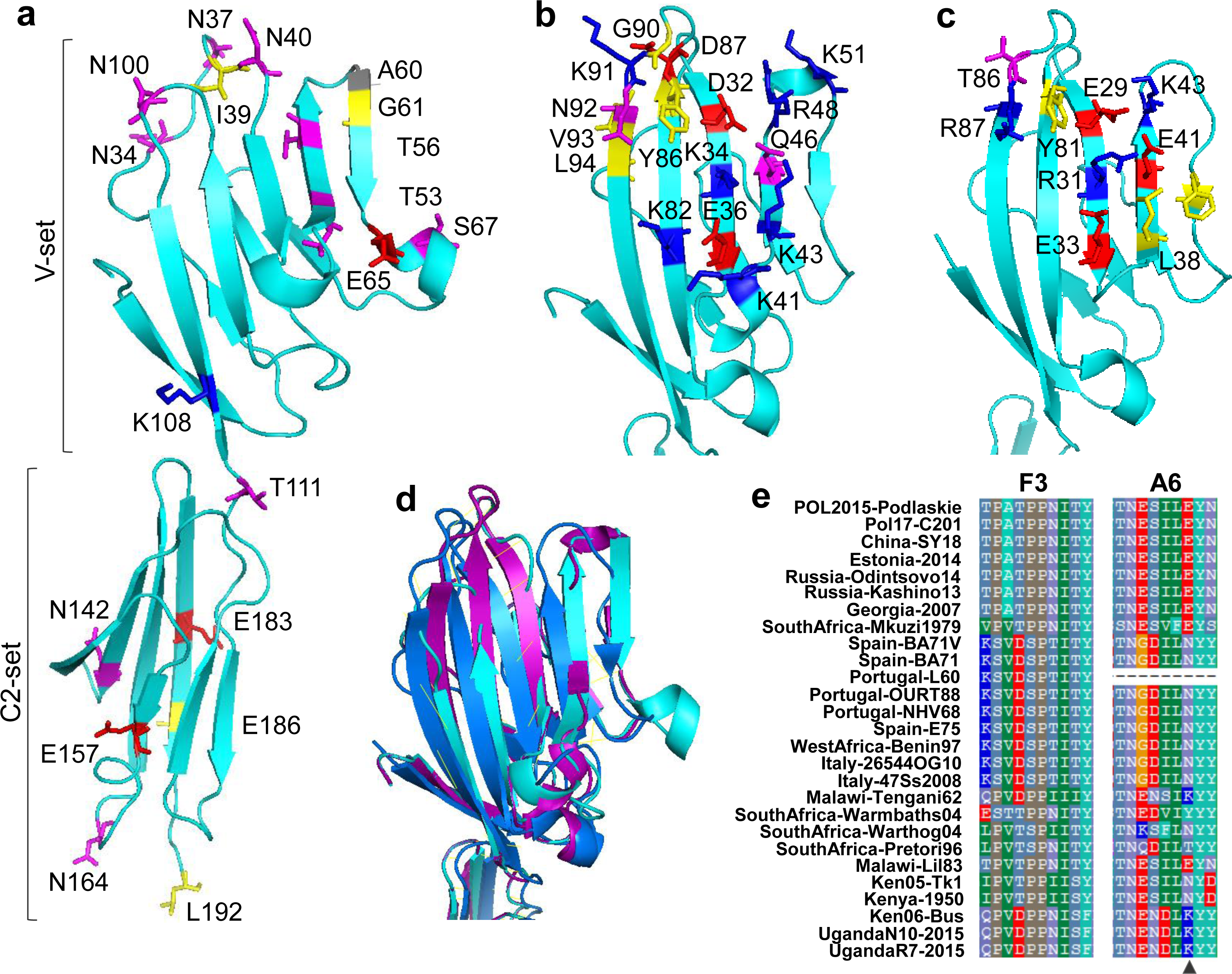
The structural mapping of the positively selected sites of EP402R and comparison with key sites in CD2 homologs. (a) The positively selected sites in EP402R mapped to the modeled structure of EP402R. Both C-set and V-set domain are shown. (b) The ligand-binding sites of human CD2 mapped to the V-set domain in the structure (PDB ID: 1hnf). (c) The ligand-binding sites of rat CD2 mapped to the V-set domain in the structure (PDB ID: 1hng). The sites are shown as colored sticks with positive-charged residues in blue, negative-charged residues in red, polar residues in magenta, and hydrophobic residues in yellow. (d) Superposition of the V-set domain of the structure of EP402R, human CD2, and rat CD2. Three proteins share a similar V-set domain structure forming a globular fold with two β-sheets. (e) Two known epitopes F3 and A6 in EP402R showing high divergence among ASFV strains. The positively selected site E157 in A6 is indicated in black triangle. The strain Portugal-L60 has a deletion at the location of A6. The truncation of EP402R by deleted nucleotides in Portugal-OURT88 and Portugal-NHV68 was recovered to obtain the normally translated epitope sequences.

The sites under positive diversifying selection have critical implications for vaccine cross-protection from heterologous viral strains when the subunits containing those sites are used as vaccines. Indeed, one of the positively selected sites E157 is located within the cytotoxic T-cell epitope A6 previously identified (Argilaguet et al., 2012). The positive diversifying selection on the site E157 and the high variability of the epitope motifs among ASFV strains provide at least partial molecular etiology of the serotype-specific T-cell response against DNA vaccines containing the epitopes in EP402R (Fig. 3e). Given the frequent occurrence of positive diversifying selection in a broad set of genes, full evaluation of the sequence variability of the target genes in designing vaccines is warranted.

A recent study of EP402R and EP153R also investigated the high sequence variability and positively selected sites in the two proteins in detail, but detected less sites under positive selection (Nefedeva et al., 2020). However, the recombination, as demonstrated in that study, might provide an alternative manifestation of positive selection exerted on ASFV, although the explicit quantification of their link might pose a new challenge.

In addition to the divergent proteins, four highly conserved structural proteins (J5R/H108R, P11.5/A137R, P10/K78R, and pp220/CP2475L, in Fig. 2c) were also found to possess positively selected sites, which have not been shown to be involved in host interactions experimentally. J5R/H108R is a transmembrane protein at the inner envelope and P10 is a DNA-binding protein in the viral nucleoid. The positive selection of the sites in these structural proteins may represent the evolutionary adaptation of ASFV for successful colonization and survival in the host niches. Another two proteins with unknown functions (MGF300-4L and B475L, in Fig. 2c), have the positively selected sites distributed across a large proportion of the gene regions. The two proteins are unique in that they exhibit high propensity for forming helices through the whole gene region. In spite of being unable to obtain confidently a tertiary structure model for the two proteins, we predicted the secondary structure of MGF300-4L and B475L using PSIPRED (Buchan and Jones, 2019). It shows that the two proteins predominantly comprise tandem α-helices. The tandem α-helix structural units have been shown to be able to stack side-by-side arranged in a specific three-dimensional conformation to create protein-binding interfaces and are commonly found in binding proteins (Groves and Barford, 1999). The presence of the tandem α-helices in the two proteins MGF300-4L and B475L indicates their possible roles in protein-protein interactions (Fig. S3).

### Identification of selective sweeps in the ASFV genomes

A selective sweep is a process where a beneficial allelic change sweeps through the population and becomes fixed in a specific population, and the nearby linked sites will hitchhike together and also become fixed. The process leads to reduced within-population genetic diversity and increased between-population differentiation in the sweeping region. Such selective sweeps allow for rapid adaptation and accelerated evolution, and are good indicators for host-pathogen interaction and adaptive evolution (Stephan, 2019). The unique mechanism of selective sweeps in causing genetic changes makes it inappropriate to detect them using the dN/dS-based method. Therefore, we detect the regions of clustered SNPs with gamma distribution, which is characteristic of SNPs under selective sweep (See Materials and Methods). We at first identified 6,054 SNPs associated with between-population subdivision and within-population homogeneity for the clade α and β (Fig. 4a). Those SNPs were subsequently subject to detection of selective sweep. A total of 578 clusters of SNPs were identified encompassing 4,741 SNPs or 2,139 non-synonymous SNPs (Supplementary file 3). That is corresponding to 26% of the total SNPs or 35% of the total non-synonymous SNPs, indicating that a high proportion of the genetic variations among the ASFV population have been likely to be introduced *via* selective sweep. Among them, 32 regions from 25 genes show high significance in the signatures of selective sweep (Fig. 4b,c and Table 2).

**Table 2.** Gene regions with significant selective sweep.

| Genomic location |  | # of SNPs | Sweep length | Gene location |  |  | *p*-value corrected | Function |
| --- | --- | --- | --- | --- | --- | --- | --- | --- |
| Start | End |  |  | Gene | Start | End |  |  |
| Genes known to be involved in host cell interaction |  |  |  |  |  |  |  |  |
| 176588 | 177082 | 56 | 494 | MGF360-16R | -1 | 493 | <1E-20 | Multigene family 360 |
| 42644 | 42992 | 55 | 349 | MGF505-9R | 12 | 360 | <1E-20 | Multigene family 505 |
| 43166 | 43394 | 28 | 229 | MGF505-9R | 534 | 762 | <1E-20 | Multigene family 505 |
| 43580 | 43775 | 24 | 196 | MGF505-9R | 948 | 1143 | <1E-20 | Multigene family 505 |
| 44899 | 45466 | 50 | 568 | MGF505-10R | 336 | 903 | <1E-20 | Multigene family 505 |
| 37872 | 38331 | 46 | 460 | MGF505-5R | 528 | 987 | <1E-20 | Multigene family 505 |
| 38379 | 38643 | 29 | 265 | MGF505-5R | 1035 | 1299 | <1E-20 | Multigene family 505 |
| 37356 | 37560 | 25 | 205 | MGF505-5R | 12 | 216 | <1E-20 | Multigene family 505 |
| 49855 | 50151 | 30 | 297 | MGF360-15R | 490 | 786 | <1E-20 | Multigene family 360 |
| 178236 | 178537 | 29 | 302 | MGF505-11L | 819 | 1120 | <1E-20 | Multigene family 505 |
| 36697 | 36997 | 25 | 301 | MGF505-4R | 902 | 1202 | 0.0150 | Multigene family 505 |
| 37041 | 37193 | 17 | 153 | MGF505-4R | 1246 | 1398 | 0.0087 | Multigene family 505 |
| 23397 | 23606 | 20 | 210 | MGF360-8L | 396 | 605 | 0.0145 | Multigene family 360 |
| 173990 | 174197 | 21 | 208 | I215L | 234 | 441 | 0.0040 | Ubiquitin conjugating enzyme |
| 46308 | 46684 | 29 | 377 | A224L | 300 | 676 | 0.0145 | IAP apoptosis inhibitor |
| Novel candidate genes |  |  |  |  |  |  |  |  |
| 150420 | 150855 | 46 | 436 | P1192R | 2890 | 3325 | <1E-20 | DNA topoisomerase type II |
| 150185 | 150371 | 18 | 187 | P1192R | 2655 | 2841 | 0.0318 | DNA topoisomerase type II |
| 22021 | 22360 | 35 | 340 | MGF300-4L | 570 | 909 | <1E-20 | Multigene family 300 |
| 48674 | 49031 | 34 | 358 | A151R | 24 | 381 | <1E-20 | Involved in redox pathway |
| 58166 | 58389 | 32 | 224 | F778R | 1167 | 1390 | <1E-20 | Ribonucleotide reductase |
| 175802 | 176124 | 31 | 323 | DP238L | 290 | 612 | <1E-20 | Uncharacterized protein |
| 156642 | 156932 | 30 | 291 | R298L | 22 | 312 | <1E-20 | Serine protein kinase |
| 63391 | 63588 | 27 | 198 | K205R | 199 | 396 | <1E-20 | Uncharacterized protein |
| 119386 | 119642 | 27 | 257 | CP2475L | 5049 | 5305 | <1E-20 | Structural polyprotein precursor |
| 160977 | 161318 | 31 | 342 | QP383R | 453 | 794 | 0.0006 | Nif S-like protein |
| 161389 | 161625 | 23 | 237 | QP383R | 865 | 1101 | 0.0029 | Nif S-like protein |
| 62145 | 62415 | 25 | 271 | F1055L | 585 | 855 | 0.0029 | Helicase superfamily II |
| 165252 | 165489 | 23 | 238 | E146L | 120 | 357 | 0.0029 | Uncharacterized protein |
| 170094 | 170377 | 24 | 284 | I267L | 68 | 351 | 0.0168 | RING finger containing protein |
| 127731 | 127897 | 20 | 167 | CP312R | 447 | 613 | 0.0006 | Uncharacterized protein |
| 67870 | 68014 | 19 | 145 | EP1242L | 2229 | 2373 | <1E-20 | RNA polymerase subunit 2 |
| 47935 | 48092 | 18 | 158 | A240L | 273 | 430 | 0.0035 | Thymidylate kinase |

Note: The significant sweeping regions should satisfy two thresholds: multiple testing corrected *p*-value $\leq 0.05$ and the number of SNPs $\geq 18$ in each region. The multiple testing corrected *p*-value was determined using the Bonferroni procedure.

**Fig. 4.**
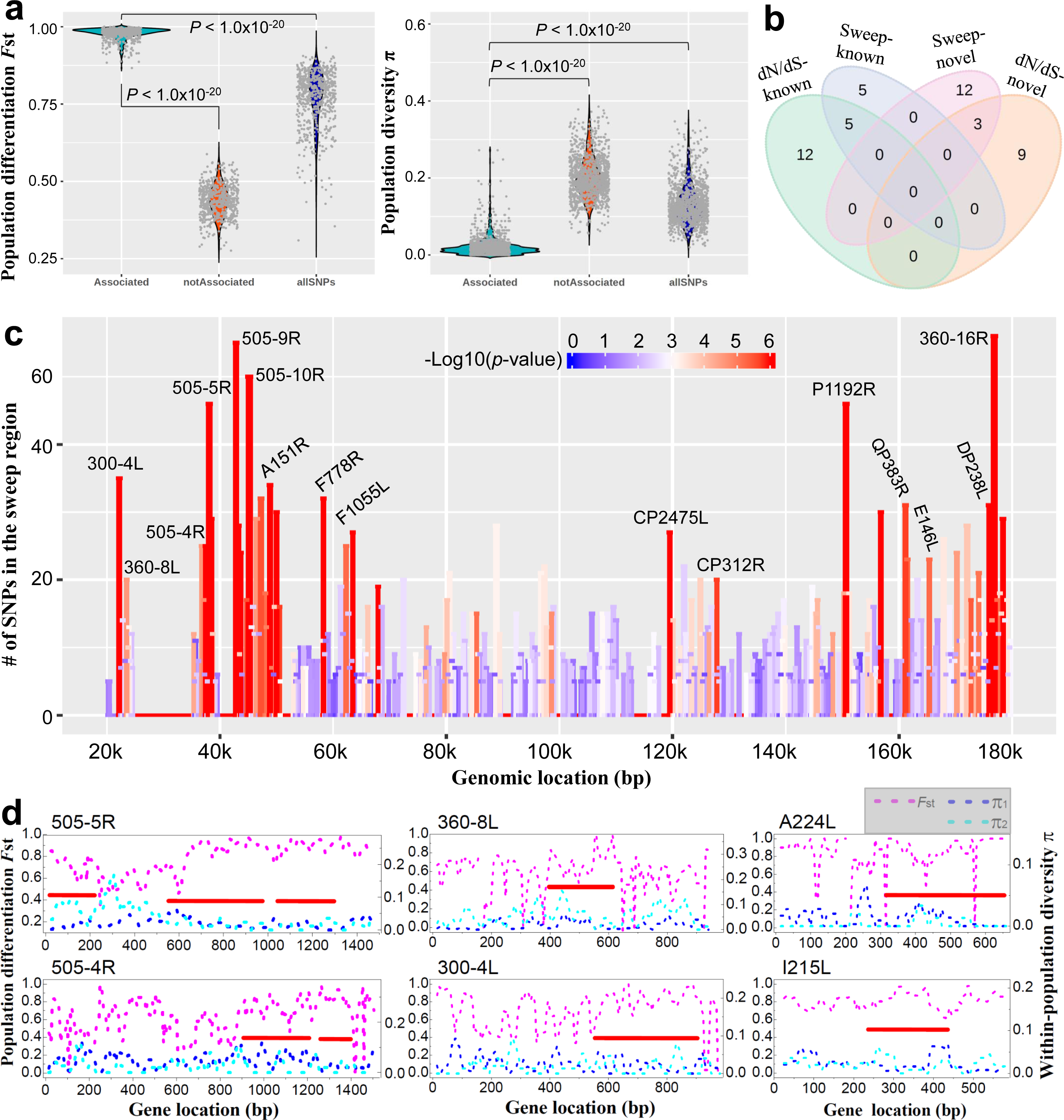
Genomic distribution and genetic properties of genes with signatures of selective sweep. (a) Distribution of population differentiation *F*_st_ and diversity π of a series of 100-loci sliding windows from three groups of SNPs: associated with between-population subdivision, not associated with between-population subdivision, and all detected SNPs. The between-group differences were evaluated using wilcoxon rank sum test and the *p*-values were indicated for the comparison between associated SNPs and the other two groups. (b) Venn diagram of number of genes with putative diversifying selection and selective sweep. (c) Significance of regions with signatures of selective sweep as shown with gradient colors. The height of bars shows the number of SNPs in the sweeping regions and the width shows the spanning length of the sweeping regions. (d) Between-population differentiation *F*_st_ (in magenta) and within-population diversity π (in blue for the clade α and cyan for the clade β) of six representative genes containing regions with putative selective sweep as shown with red bars. Only the sweeping regions longer than 135bp and Bonferroni-corrected *p*-value ≤ 0.05 were considered significant and indicated. The regions show higher between-population differentiation and reduced within-population diversity in comparison with the nearby regions. The scale for the between-population differentiation is shown on the left axis and the within-population diversity on the right axis.

The gene regions with significant selective sweep exhibit higher population differentiation and reduced sequence diversity as shown in the key signature genes (Fig. 4d). Among them are a series of known gene factors involved in host cell interactions, including MGF505, MGF360 and I215L, which also harbor sites under positive diversifying selection. Those gene factors exhibit genetic signatures of both diversifying selection and selective sweep (Fig. 4d and Fig. 2b). Noteworthy are the 15 novel candidate genes showing strong signatures of selective sweep (Table 2). A large proportion of them (60%) are involved in key cellular functions, such as replication, repair, transcription, and metabolism (Table 2).

We notice that four of the novel candidates (A151R, F1055L, CP312R, and E146L) have been previously demonstrated to induce immune responses in swine following ASFV challenge (Jancovich et al., 2018; Netherton et al., 2019). Therefore, we proceed to characterize the shared genetic properties of the candidate genes and compare with that of known genes inducing immune responses or involved in host cell interaction.

### Sequence variability of the candidate genes with diversifying selection or selective sweep

We ascertain the genetic properties of the genes with positive diversifying selection or selective sweep by calculating population prevalence frequencies and pair-wise amino acid divergence of the genes and doing comparison with three gene categories cataloged from other studies: (i) the non-antigenic conserved structural proteins without positive selection (Alejo et al., 2018), (ii) the antigen proteins eliciting immunological responses in immunoassay experiments (Jancovich et al., 2018; Lopera-Madrid et al., 2017; Netherton et al., 2019), (iii) the proteins previously shown to be involved in host cell interactions (Dixon et al., 2013; Dixon et al., 2019) (Fig. 5 and Table S7). A non-uniform population prevalence and higher level of sequence variability are observed in the candidate genes under putative positive diversifying selection in comparison with the category of (i) conserved structural proteins and (ii) antigenic proteins, but not with the gene category (iii) involved in host cell interactions (two-sided Mann-Whitney U-test, Fig. 5a,d,i). The overall high divergence in amino acid sequences coupled with the significant positive diversifying selection of those genes suggests that they have mutated frequently during evolution. In contrast, the candidate genes with signatures of selective sweep are relatively more conserved and present a comparable level of sequence variability with that of conserved structural proteins and the known antigenic proteins, supporting their potentiality as generalized immunogenic targets (Fig. 5b,e,i).

**Fig. 5.**
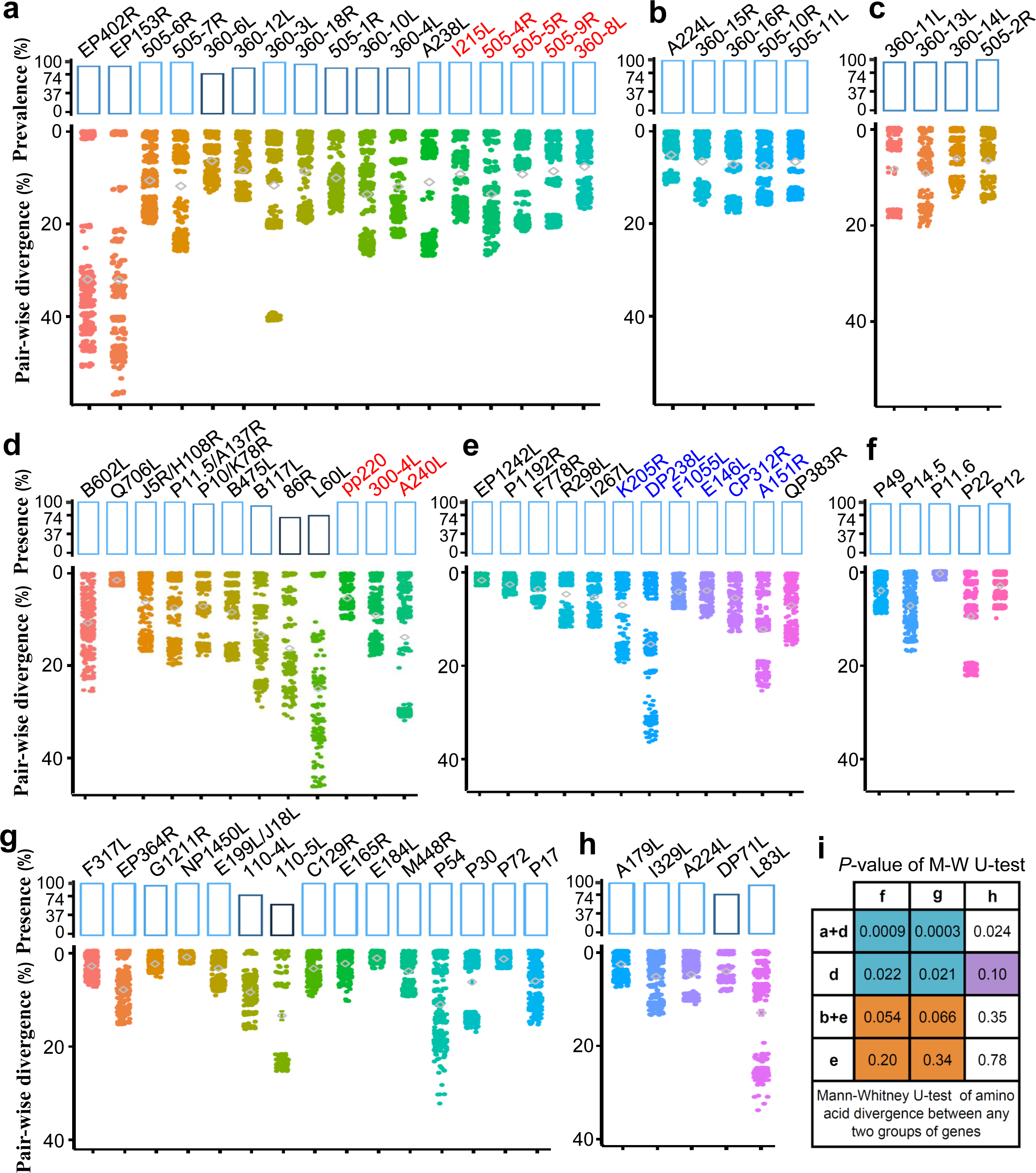
Presence frequencies and sequence divergence of the genes with signatures of diversifying selection and those with selective sweep. (a) The genes with signals of diversifying selection in this study and known to be involved in host interactions. (b) The genes with selective sweep in this study and known to be involved in host interactions. (c) The genes lost in avirulent strains without significant diversifying selection or selective sweep. (d) The novel candidate genes with diversifying selection signals. (e) The novel candidate genes with selective sweep signals (f) The non-antigenic conserved structural proteins. (g) The antigen proteins eliciting immunological responses in immunoassay experiments. (h) The genes known to be involved in interactions with host cell components. (i) Mann-Whitney U-test of amino acid divergence between any two groups of genes above. For each gene, the mean amino acid divergence among the ASFV strains was used as the proxy for the test. The presence frequency was calculated as the percentage of presence of each gene within the 27 non-redundant ASFV strains and represented as colored bars. The sequence divergence was evaluated as pair-wise amino acid differences displayed as jitter plots. The average of pair-wise divergence for each gene is indicated with grey diamond. The names of MGF genes ignore “MGF” for figure compactness.

### Genetic diversity and divergent selections among paralogous gene members of MGF360/505

Given that a large number of MGF genes have been identified to be genetically diverse with significant signatures of positive selection, a natural question is: how about the breath of genetic diversity and selection pressures among the paralogous members of MGF and which regions are responsible for the genetic and functional diversity? We examine the genetic diversity of MGF genes by evaluating the differential selection between paralogous genes/branches of the two families MGF360 or MGF505. We first constructed the phylogenetic structures of all orthologous and paralogous members of MGF360 and MGF505, respectively (Fig. 6a,c and Fig. S4), and then chose the phylogenetically close pairs of genes/branches to perform the likelihood ratio test of divergent selection. The test identified 10 and 9 pairs showing divergent selection on an average of 8.3% and 9.6% of the sites among MGF360 and MGF505, respectively (*p*-value ≤ 0.05, Chi-squared test) (Fig. 6a,c and Table S8). The divergent selection clearly indicates the distinct evolutionary forces exerted on the array of paralogs of MGF, thus forming a genetic pool for functional diversification. The functional diversification is further supported by the divergent regulation patterns across the paralogous members of MGF (Fig. 6b,d). The regulatory divergence is manifested qualitatively in the distinct promoter motifs and their distances to the translation start site (TSS) among paralogous members of MGF. Further profiling the promoter regions 55 nucleotides upstream TSS of MGF genes shows that the promoter divergence is correlated with the evolutionary distances between paralogs of MGF (Fig. S5). The regulatory divergence in the promoter regions, coupled with the differentiated selection pressures between paralogous pairs of MGF360 and MGF505 constitutes important genetic basis for functional diversification of MGF genes, providing a wide spectrum of specificity in host tropism and adaptation. Interestingly, we found that the consensus motif patterns we obtained specifically for MGF360/MGF505 are very similar to that profiled experimentally in a recent study for a set of early transcribed genes including MGF genes in the ASFV strain Spain-BA71V (Cackett et al., 2020). The consistency between the two studies provides further support for our results.

**Fig. 6.**
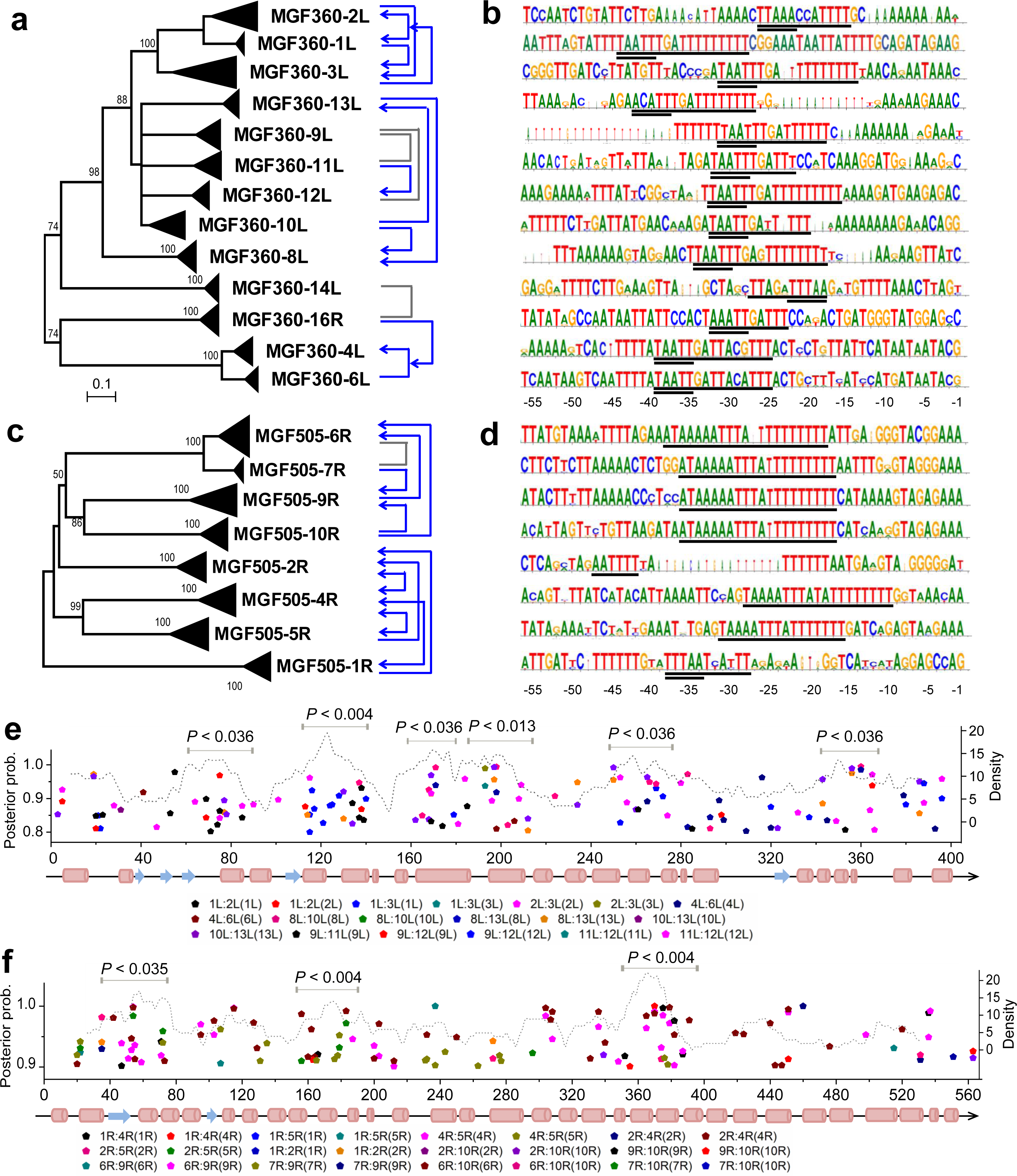
Genetic diversity among paralogs of MGF360 and MGF505. (a,c) Divergent selection between paralogous pairs of genes/branches of MGF360 and MGF505 mapping to the phylogenetic structure. The phylogenetic trees were inferred using Neighbor-Joining method with 1000 bootstraps. The branches containing orthologous members of each paralog are collapsed indicated with triangle. The exceptions are three isolates of MGF360-1L (Kenya-1950, Ken05-Tk1, and Spain-E75), which cluster together with MGF360-2L, and five isolates of MGF505-7R (Malawi-Lil83, Kenya-1950, Ken05-Tk1, Ken06-Bus, and UgandaN10-2015), which cluster together with MGF505-6R. The pairs of genes/branches used for LRTs are connected by frame lines with blue arrows indicating the gene/branch under positive selection at a fraction of sites and grey lines indicating no significant positive selection in either of the gene/branch. (b,d) Divergent promoter regions from -55 to -1 upstream translational start sites of MGF360 and MGF505. The sequences with common signatures are highlighted with underline and the potential 5-nucleotide promoter motifs with double underline. (e,f) Distribution and enrichment of sites under divergent selection between paralogous pairs of genes/branches of MGF360 (e) and MGF505 (f). Only the sites with a posterior probability ≥ 0.8 in MGF360 and ≥ 0.9 in MGF505 are shown (colored pentagons). Either of the partners in the pairs was treated as foreground in LRTs (indicated in the parentheses). The sites are mapped to the predicted secondary structure of MGF360 and MGF505, respectively (cylinders for α-helices, arrows for β-strands, and lines for coiled loops. A 25-codon sliding window plot of the site density is shown as dotted grey lines. The *p*-value of enrichment was calculated with the Hypergeometric test for each 25-codon window and the consecutive windows with *p*-value ≤ 0.05 were merged to a single region indicated with horizontal bars.

To unveil the genetic properties of the gene regions under divergent selection, we identified the sites under putative divergent selection between the paired genes/branches of MGF360/MGF505, and quantified the site distribution along the predicted secondary structure of MGF360/MGF505, respectively (Fig. 6e,f and Supplementary file 4). Interestingly, the sites exhibit quasi-periodic distribution and are enriched periodically in a few patches of length ∼ 30 residues (*p*-value ≤ 0.05, Hypergeometric test). This average length of enrichment is close to the length of the ankyrin repeat (Mosavi et al., 2004), which is believed to be the building blocks of the MGF protein structures. Actually, the predicted secondary structures of MGF360 and MGF505 display signatures of tandem ankyrin repeats, each consisting of a helix-loop-helix motif followed by another loop region. Protein domains containing tandem ankyrin repeats usually fold into a conserved tertiary concave/convex structure mediating protein-protein interactions. The surface recognition residues are highly variable, affording specific interactions with a broad range of host targets (Mosavi et al., 2004). Ankyrin repeats have been described to be the major functional units in host range factors in several poxvirus species (Bradley and Terajima, 2005; Herbert et al., 2015; Li et al., 2010). Here in the absence of the protein structure of MGF proteins, we demonstrated that the periodic patches of residues in ankyrin repeats exhibit differentiated evolutionary selection among paralogous members, thereby representing the motifs facilitating genetic and functional diversity of MGF in the multifaceted interactions with host cells. Further studies are required to ascertain the role of the motifs in host interactions.

## Discussion

In our pursuit of characterizing the variation landscape of ASFV genomes and unraveling a comprehensive set of candidate genes with positive selection signatures and relevance to host adaptation and interaction, we identified 29 candidate genes with positive diversifying selection and 25 with selective sweep. Among them, eight show signatures of both kinds of selection and 24 are novel candidates that so far, have not been reported to be associated with host interactions. The genes showing selection signatures are widely distributed across the genome, highlighting adaptive evolution at multiple genomic regions of ASFV during the interactions with hosts. We summarize and present the candidate genes in a unified scheme of interactions between ASFV and hosts in a framework of the virus life cycles and host defense processes (Fig. 7) (Rodriguez and Salas, 2013).

**Fig. 7.**
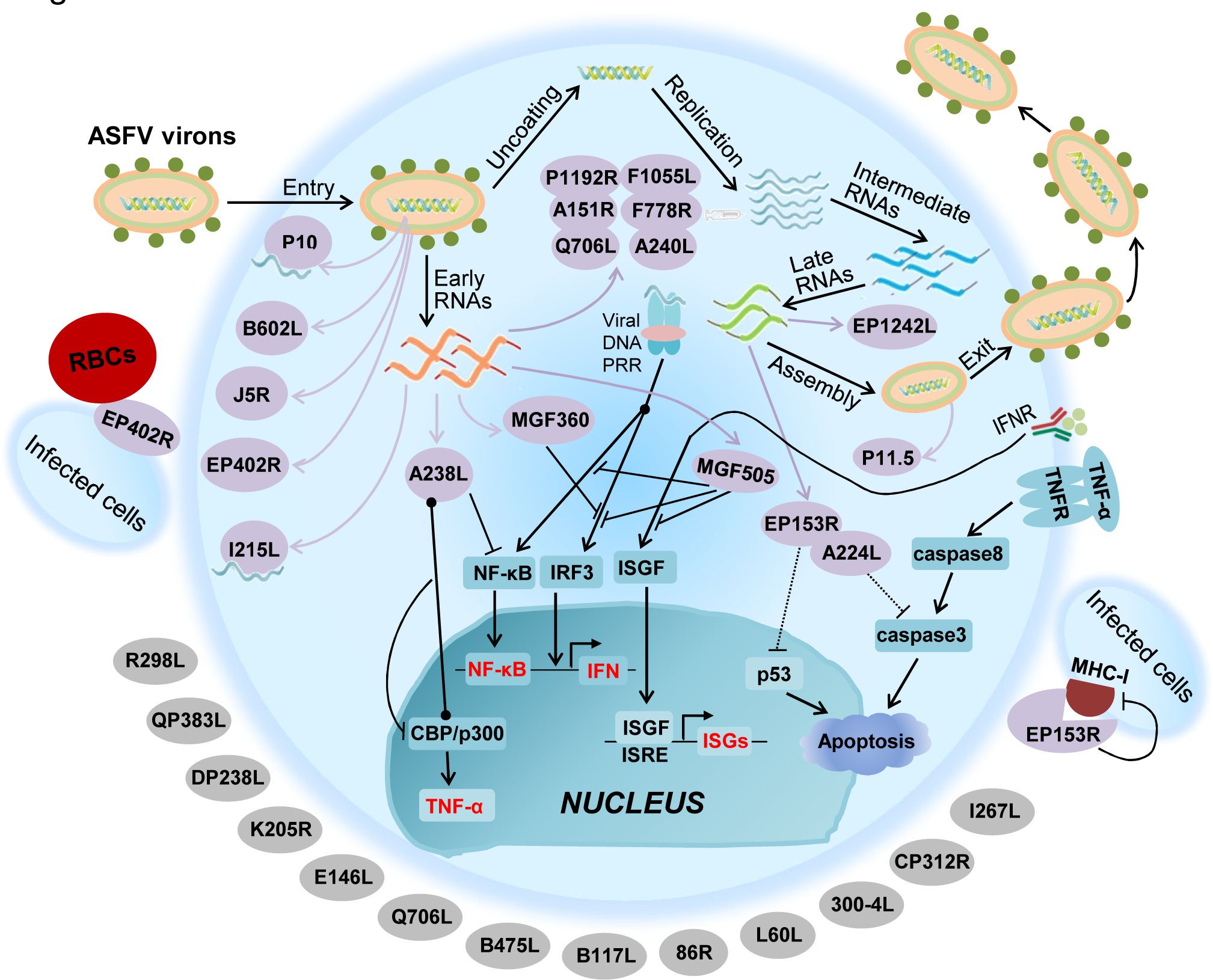
The integrated scheme of interactions between ASFV genes with signatures of diversifying selection/selective sweep and host components. The interactions are depicted in the framework of the virus life cycles and host defense processes. The ASFV-encoded proteins are associated with different parts of the viral particle or released at different stages of the infection cycle (purple ovals). They interact with host cells *via* DNA-binding, surface adhesion, inhibition, or activation. The host cell is bounded with membrane indicated with the round soft edge. Host-encoded proteins are shown as aqua squares. ASFV-encoded proteins with unknown function or expression time are shown as grey ovals outside of the membrane. Not all members of MGF360 or MGF505 are involved in the interactions. Key host molecules affected by ASFV, such as NF-κB, IFN, TNF-α, and ISGs are shown in red. Other abbreviations: TNFR: TNF receptor; IFNR: IFN receptor; Viral DNA PRR: viral DNA pattern recognition receptor; ISGF: IFN-stimulated gene factor; ISGs: IFN-stimulated genes; ISRE: IFN-stimulated response elements; RBCs: red blood cells.

The proteins in the scheme include those known to be relevant to host immune evasion, such as EP402R for surface adherence of infected cell (Borca et al., 1998), EP153R for inhibition of MHC expression and host cell apoptosis (Alejo et al., 2018; Hurtado et al., 2011), A238L for production impairment of immune regulator NF-κB and cytokines TNF-α (Powell et al., 1996), and multiple MGF genes for modulation of interferon (IFN) response (Afonso et al., 2004; Correia et al., 2013).

The scheme also contains the proteins critical for the virus life cycles facilitating successful entry and proliferation in host cells, such as the structural proteins pp220, J5R, P11.5, P10, and B602L localizing at distinct layers of the viral particles for virus entry and assembly (Alejo et al., 2018), the basic enzymes P1192R, F1055L, F778R, A240L and EP1242L involved in replication, repair and transcription in host cytoplasm (Dixon et al., 2013). The key roles played by the proteins and the relatively high conservation make them promising candidates for vaccines with cross-activity.

The cellular processes the candidate genes are involved in, provide a variety of sources of selective pressures acting at multiple stages of the infection cycles for ASFV to evolve and adapt. In this regard, these genes may constitute an important part of the genetic factors of ASFV in circumventing host defense systems and enhancing fitness in a specific manner.

Our data reveals that the adaptive evolution of ASFV has been shaped by both positive diversifying selection and selective sweep. The results show that the genes with diversifying selection exhibit a higher level of sequence variability than those with selective sweep and provide important implications for vaccine design. The most prominent are EP402R, EP153R and MGF360/MGF505 with the highest genetic variability, the only known proteins so far shown to be both virulence determinants and immunogenic targets (Boinas et al., 2004; Burmakina et al., 2016). However, the high sequence diversity of EP402R/EP153R and mosaic presence pattern of MGF360/MGF505 among the ASFV population make it difficult for them to achieve desirable cross-protection (Malogolovkin et al., 2015). The dual role of EP402R, EP153R and MGF360/MGF505, as both potential virulence determinants and immunogenic proteins, may also introduce confounding factors in designing live-attenuated virus vaccines (LAVs). Recently, as an encouraging example, elimination of EP402R from the virulent BA71 to obtain the LAV strain BA71ΔCD2, protected pigs against homologous and heterologous virus challenges (Monteagudo et al., 2017). Similarly, ASFV-Georgia-ΔMGF, a LAV strain lacking a series of MGF genes, protected animals against homologous challenges (Donnell et al., 2015). Unfortunately, sequential deletion of multiple genes provoked in occasion the loss of protection due to excessive attenuation (O’Donnell et al., 2016). It is worth mentioning that the role of EP402R as a virulence factor of ASFV has not yet been explicitly determined due to differential virulence outcomes from disruption of EP402R in distinct isolates. A few studies have shown that abrogation of EP402R function does not significantly alter the virulence of the mutants (Borca et al., 1998; Borca et al., 2020). The isolate-dependent functional effect of EP402R will pose additional challenges for designing LAVs.

The divergent selection between paralogous genes of MGF360/MGF505 further complicates the vaccine design. We identified differentiated selection pressures and regulation patterns between paralogs of MGF360/MGF505 conferring genetic diversity and functional diversification. The possible scenario is that the antigenic activities and expression levels of paralogs of MGF360/MGF505 are strain-specific and/or host-dependent. This scenario provides a rationale for the observations that variable deletion patterns and expression profiles of paralogous members of MGF have been resulted from different adaptation processes or have induced distinct viral growth outcomes in host niches (Krug et al., 2015; Rodríguez et al., 2015). Up to now, the precise connections between the MGF genes and physiological conditions are still largely unknown. Optimal choices of paralogous MGF genes and gene regions remain to be tested when they are used as immunogenic targets. The specific sites under divergent selection we dissected in MGF360/MGF505 provide important information in aiding for the tests.

Compared to the high divergence of the candidate genes with diversifying selection, the genes with selective sweep display a relatively low level of within-population diversity at sweeping regions and a high degree of average conservation. Many of them (60% of the novel candidates) are involved in the critical events in the life cycles of ASFV infections, such as replication, repair and transcription. Interestingly, an evolutionary study of the influenza A virus H3N2 showed that the emergent severe seasonal flu in 2004/2005 was correlated with mutations in the key ribonucleoprotein (RNP) complex acquired by a circulating lineage *via* selective sweep and the lineage was demonstrated to induce elevated replicative fitness and more severe clinical diseases (Memoli et al., 2009). We argue that the genes with selective sweep are important contributing factors for the rapid adaptation and enhanced fitness of the ASFV population circulating in specific areas. The relatively high conservation and critical roles of the genes make them promising candidates for vaccine molecules or drug targets.

We highlight the importance of our findings in the two following two aspects. (1) Our data provides novel insights into the adaptation and fitness of ASFV. The multifaceted genetic characteristics of ASFV genes imply that the virus may have utilizing multiple mechanisms (such as genetic diversification, selective sweep and divergent selection) and pertinent genetic factors for successful replication, adaption, and persistence during interaction with continuously changing host environments, including warthogs, ticks, and domestic pigs. The plethora of variable genetic factors may act as a genetic pool for adaptation to new hosts by functional diversification or immune escape, though the alternative new hosts beyond the currently known have not yet been reported. (2) The candidate genes we identified in the study could serve as valuable targets for vaccine molecules or therapeutic agents, and the sites with signatures of positive selection will be valuable for precise design and engineering. We also understand that the methods we used for identifying selection are not perfect and the genetic variability might have been underestimated due to the limited size of the ASFV population or the conservation of the selection analysis methods. We believe that the availability of more genomic information in the future will be of great help for overcoming the limitation.

## Data availability

The multiple sequence alignments used for selection analysis and supplementary files are available through the links: https://figshare.com/projects/ASFV_alignment/82718 and https://figshare.com/projects/ASFV_supplementary_files/90335, respectively under the MIT license.

## Acknowledgements

The work was supported by the National Key Research and Development Program of China (2018YFC0840401).

## Ethics statement

The authors confirm that the ethical policies of the journal, as noted on the journal’s author guidelines page, have been adhered to. No ethical approval was required as this is a meta-analysis article.

---

**Table S1.** Information of ASFV isolates with known genomic sequences.

| **Genome accession** | **Virulence** | **Strain name** | **Name tag in this study** | **Isolation location** |
| --- | --- | --- | --- | --- |
| ASU18466 | Low | BA71V | Spain-BA71V | Spain: Badajoz |
| KP055815 | High | BA71 | Spain-BA71V | Spain: Badajoz |
| KM262844 | High | L60 | Portugal-L60 | Portugal |
| KM262845 | Low | NHV | Portugal-NHV68 | Portugal |
| AY261360 | High | Kenya 1950 | Kenya-1950 | Kenya |
| KM111294 | Moderate | Ken05/Tk1 | Ken05-Tk1 | Kenya central |
| KM111295 | High | Ken06.Bus | Ken06-Bus | Kenya eastern |
| AY261366 | Unknown | Warthog | Namibia-Warthog04 | Namibia |
| AY261365 | Unknown | Warmbaths | SouthAfrica-WarmBaths04 | South Africa: Warmbaths |
| AY261364 | High | Tengani 62 | Malawi-Tengani62 | Malawi: Tengani, Nsanje District |
| AY261363 | High | Pretorisuskop/96/4 | SouthAfrica-Pretori96 | South Africa: Kruger National Park |
| AY261362 | Unknown | Mkuzi 1979 | SouthAfrica-Mkuzi1979 | South Africa: Mkuzi Game Reserve |
| AY261361 | High | Malawi Lil-20/1 | Malawi-Lil83 | Malawi: Chalaswa |
| AM712239 | High | Benin 97/1 | WestAfrica-Benin97 | West Africa: Benin |
| AM712240 | Low | OURT 88/3 | Portugal-OURT88 | Portugal |
| FN557520 | High | E75 | Spain-E75 | Spain: Lerida |
| FR682468 | High | Georgia 2007/1 | Georgia-2007 | Georgia |
| KX354450 | High | 47/Ss/2008 | Italy-47Ss2008 | Italy: Province of Sassari, Sardinia |
| KM102979 | High | 26544/OG10 | Italy-26544OG10 | Italy |
| KJ747406 | Medium | Kashino 04/13 | Russia-Kashino13 | Russia |
| LS478113 | Unknown | Estonia 2014 | Estonia-2014 | Estonia |
| KP843857 | High | Odintsovo_02/14 | Russia-Odintsovo14 | Russia |
| MH025916 | Unknown | R8 | UgandaR8-2015 | Uganda: Tororo district |
| MH025917 | Unknown | R7 | UgandaR7-2015 | Uganda: Tororo district |
| MH025918 | Unknown | R25 | UgandaR25-2015 | Uganda: Tororo district |
| MH025919 | Unknown | N10 | UgandaN10-2015 | Uganda: Tororo district |
| MH025920 | Unknown | R35 | UgandaR35-2015 | Uganda: Tororo district |
| MH681419 | High | POL/2015/Podlaskie | POL2015-Podlaskie | Poland |
| MG939583 | Unknown | Pol16_20186_o7 | Pol16-o23 | Poland |
| MG939584 | Unknown | Pol16_20538_o9 | Pol16-o9 | Poland |
| MG939585 | Unknown | Pol16_20540_o10 | Pol16-o10 | Poland |
| MG939586 | Unknown | Pol16_29413_o23 | Pol16-o23 | Poland |
| MG939587 | Unknown | Pol17_03029_C201 | Pol17-C201 | Poland |
| MG939588 | Unknown | Pol17_04461_C210 | Pol17-C210 | Poland |
| MG939589 | Unknown | Pol17_05838_C220 | Pol17-C220 | Poland |
| MH766894 | Unknown | ASFV-SY18 | China-SY18 | China |

Note: The strains highlighted in grey are excluded from analysis due to the high similarity with other strains of the same subtypes. Therefore, a total of 27 non-redundant strains were used for this study.

**Table S2.** List of unique non-synonymous mutations in the strains with low virulence.

| Gene | AA change | Nucleotide change | Gene function |
| --- | --- | --- | --- |
| F1055L | Met114Ile | 342C>T | Helicase superfamily II |
| C147L | Met11Val | 33T>C | RNA polymerase subunit 6 |
| B119L | Phe18Ser | 54A>G | Component of redox pathway |
| CP204L/P30 | Thr116Ala | 348T>C | Phosphoprotein binds to ribonucleoprotein-K |
| CP312R | Pro190His | 569C>A | Hypothetical protein |
| NP868R | Asp182Gly | 546A>G | Guanylyl transferase (for mRNA modification) |
| NP868R | Gln259Arg | 777A>G |  |
| E199L/J18L | Gln61Arg | 183T>C | Transmembrane domain containing protein |
| E120R/P14.5 | His111Arg | 333A>G | DNA-binding. Required for movement of virions to plasma membrane |
| I215L | Glu95Gly | 285T>C | Ubiquitin conjugating enzyme |
| MGF505-11L | His251Arg | 753T>C | Multigene family 505 |
| MGF505-11L | Leu213Phe | 639C>G |  |
| MGF505-11L | Lys131Arg | 393T>C |  |

Note: The unique mutation is defined as those occur in the two ASFV strains with low virulence (Portugal-NHV68 and Portugal-OURT88). The positions are relative to the strain Georgia-2007.

**Table S3.** ASFV genes enriched with non-synonymous mutations.

| Gene name | Non-synonymous mutation counts | Gene length | *p*-value | *p*-value corrected | Gene function | Functional category |
| --- | --- | --- | --- | --- | --- | --- |
| MGF300-4L | 116 | 993 | <1E-20 | <1E-20 | MGF300-4L | Multigene family |
| MGF300-1L | 74 | 807 | 2.57E-09 | 2.35E-08 | MGF300-1L | Multigene family |
| MGF505-4R | 274 | 1521 | <1E-20 | <1E-20 | MGF505-4R | Multigene family |
| MGF505-5R | 186 | 1497 | <1E-20 | <1E-20 | MGF505-5R | Multigene family |
| MGF505-6R | 99 | 1578 | 0.0002 | 0.00115 | MGF505-6R | Multigene family |
| MGF505-9R | 192 | 1521 | <1E-20 | <1E-20 | MGF505-9R | Multigene family |
| MGF505-10R | 145 | 1629 | <1E-20 | <1E-20 | MGF505-10R | Multigene family |
| MGF505-11L | 128 | 1629 | 1.99E-10 | 2.13E-9 | MGF505-11L | Multigene family |
| MGF360-8L | 118 | 960 | <1E-20 | <1E-20 | MGF360-8L | Multigene family |
| MGF360-15R | 75 | 870 | 2.71E-08 | 1.92E-07 | MGF360-15R | Multigene family |
| MGF360-16R | 93 | 930 | 2.08E-13 | 2.67E-12 | MGF360-16R | Multigene family |
| A151R | 85 | 477 | <1E-20 | <1E-20 | CXXC-motif containing protein | Involved in redox pathway |
| I215L | 78 | 639 | <1E-20 | <1E-20 | Ubiquitin-conjugation enzyme | Shuttles between the nucleus and cytoplasm |
| I196L | 72 | 609 | 3.51E-14 | 4.99E-13 | Uncharacterized protein |  |
| I177L | 31 | 201 | 1.14E-09 | 1.12E-08 | Uncharacterized protein |  |
| DP238L | 68 | 717 | 2.88E-09 | 2.46E-08 | Uncharacterized protein |  |
| H240R | 68 | 726 | 4.76E-09 | 3.81E-08 | Uncharacterized protein |  |
| K205R | 59 | 618 | 2.42E-08 | 1.82E-07 | Uncharacterized protein |  |
| E183L/P54 | 49 | 555 | 3.25E-06 | 2.08E-05 | Structural protein p54 | Structural protein |
| A240L | 58 | 711 | 5.17E-06 | 5.15E-05 | Thymidylate kinase | Nucleotide metabolism |
| EP364R | 79 | 1110 | 1.94E-05 | 1.13E-04 | ERCC4 domain | DNA replication and repair |
| I267L | 61 | 840 | 9.21E-05 | 5.13E-04 | RING finger containing protein |  |
| CP312R | 65 | 924 | 1.40E-04 | 7.33E-04 | Uncharacterized protein |  |
| A137R/P11.5 | 35 | 414 | 1.80E-04 | 8.98E-04 | Structural protein P11.5 | Structural protein |
| I329L | 68 | 990 | 1.90E-04 | 9.54E-4 | Transmembrane protein | Host-cell interactions |

Note: The enrichment *p*-value for each gene was calculated with Hypergeometric test and the multiple testing correction was determined using the Benjamini-Hochberg procedure. The enrichment with corrected *p*-value < 0.001 is considered to be significant.

**Table S4.**
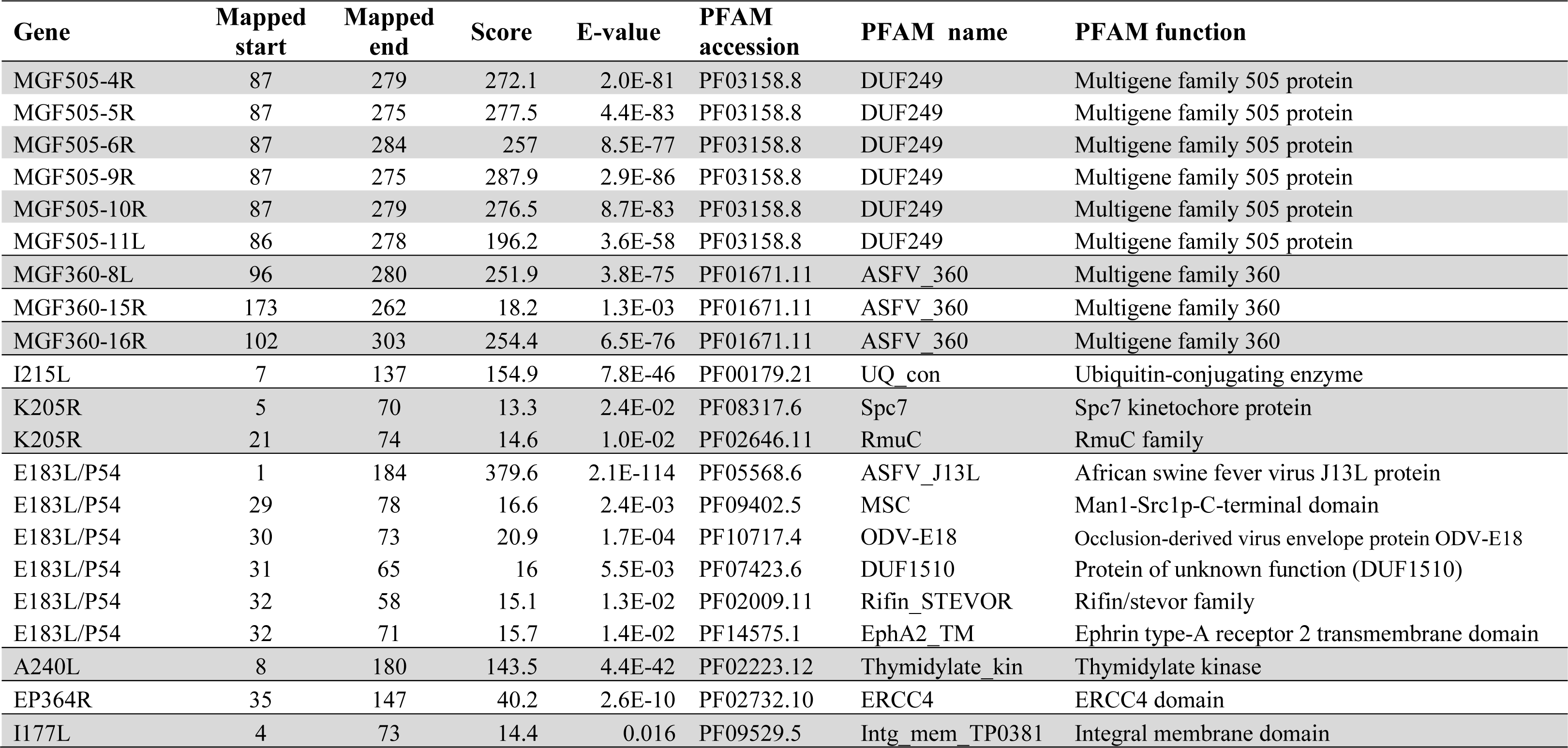
Functional domain identification of the genes enriched with non-synonymous mutations (*E*-value ≤ 0.03 or score ≥ 20).

**Table S5.** Genes with the value of dN/dS lower than the average (dN/dS < 0.1) using the Nei & Gojobori method.

| **Gene** | **dN/dS** | **Gene function** | **Functional category** |
| --- | --- | --- | --- |
| NP1450L | 0.100 | RNA polymerase subunit 1 | Transcription |
| A859L | 0.098 | Helicase superfamily II | Transcription |
| NP419L | 0.093 | DNA ligase | DNA replication |
| E165R | 0.093 | dUTPase | DNA metabolism |
| M1249L | 0.090 | Ubiquitin-like domain containing protein |  |
| D205R | 0.087 | RNA polymerase subunit 5 | Transcription |
| P1192R | 0.085 | Topoisomerase II | DNA replication |
| H359L | 0.082 | RNA polymerase subunit 3 | Transcription |
| NP868R | 0.076 | mRNA guanylyltransferase | Transcription |
| F1055L | 0.074 | Helicase superfamily II | Transcription |
| F778R | 0.072 | Ribonucleotide reductase large subunit | Nucleotide metabolism, transcription, replication and repair |
| C147L | 0.071 | RNA polymerase subunit 6 | Transcription |
| S273R | 0.069 | Ulp1 protease Family | Structural protein |
| B263R | 0.068 | TATA-box binding-like protein | Nucleotide metabolism, transcription, replication and repair |
| E184L | 0.066 | Hypothetical protein |  |
| C962R | 0.064 | Putative DNA primase | DNA replication |
| F334L | 0.061 | Ribonucleotide reductase small subunit | Nucleotide metabolism, transcription, replication and repair |
| EP424R | 0.060 | FTS J-like Methyltransferase domain containing protein | Nucleotide metabolism, transcription, replication and repair |
| B385R | 0.054 | A2L-like transcription factor | Transcription |
| B646L/P72 | 0.046 | Structural protein P72 | Structural protein |
| CP80R | 0.040 | RNA polymerase subunit 10 | Transcription |
| CP530R | 0.039 | 60 kDa polyprotein | Structural protein |
| E301R | 0.035 | Proliferating cell nuclear antigen-like protein | DNA replication |
| B125R | 0.028 | E2 early regulatory protein | Regulator of transcription and DNA replication |
| C315R | 0.025 | TFIIB like | Transcription |
| B354L | 0.022 | P-loop-containing nucleoside triphosphate hydrolases | Energy metabolism |
| EP1242L | 0.019 | RNA polymerase subunit 2 | Transcription |
| A104R/P11.6 | 0.004 | Histone-like structural protein | Structural protein |

**Table S6.** Genes with positive selection signals at a fraction of sites with . (dN/dS) >1 based on the likelihood ratio tests.

| Gene | Parameters for M2 |  | Parameters for M8 |  | M2 vs. M1 |  | M8 vs. M7 |  |
| --- | --- | --- | --- | --- | --- | --- | --- | --- |
|  |  |  |  |  | LRT<sup>a</sup> | *p*-value | LRT<sup>a</sup> | *p*-value |
| CP2475L | $p_2 = 0.013$, | $\omega_2 = 4.642$ | $p_1 = 0.018$, | $\omega = 4.001$ | 63.492 | 1.63E-14 | 78.773 | <1E-20 |
| MGF505-4R | $p_2 = 0.040$, | $\omega_2 = 5.500$ | $p_1 = 0.044$, | $\omega = 5.371$ | 43.337 | 3.89E-10 | 45.324 | 1.44E-10 |
| EP402R | $p_2 = 0.046$, | $\omega_2 = 4.297$ | $p_1 = 0.065$, | $\omega = 3.117$ | 34.179 | 3.78E-08 | 45.886 | 1.09E-10 |
| I215L | $p_2 = 0.057$, | $\omega_2 = 6.255$ | $p_1 = 0.060$, | $\omega = 6.008$ | 31.813 | 1.24E-07 | 32.542 | 8.58E-08 |
| EP153R | $p_2 = 0.104$, | $\omega_2 = 4.427$ | $p_1 = 0.113$, | $\omega = 3.639$ | 30.659 | 2.20E-07 | 30.156 | 2.83E-07 |
| MGF505-6R | $p_2 = 0.044$, | $\omega_2 = 3.617$ | $p_1 = 0.054$, | $\omega = 3.351$ | 20.746 | 3.13E-05 | 23.032 | 9.97E-06 |
| B117L | $p_2 = 0.025$, | $\omega_2 = 246.8$ | $p_1 = 0.025$, | $\omega = 232.2$ | 23.076 | 9.75E-06 | 22.893 | 1.07E-05 |
| MGF505-9R | $p_2 = 0.049$, | $\omega_2 = 4.534$ | $p_1 = 0.054$, | $\omega = 4.334$ | 22.579 | 1.25E-05 | 22.881 | 1.08E-05 |
| B602L | $p_2 = 0.046$, | $\omega_2 = 3.788$ | $p_1 = 0.051$, | $\omega = 3.665$ | 19.422 | 6.06E-06 | 21.450 | 2.20E-05 |
| 86R | $p_2 = 0.131$, | $\omega_2 = 10.373$ | $p_1 = 0.131$, | $\omega = 10.373$ | 19.944 | 4.67E-05 | 19.976 | 4.60E-05 |
| MGF505-7R | $p_2 = 0.057$, | $\omega_2 = 3.524$ | $p_1 = 0.067$, | $\omega = 3.318$ | 17.985 | 1.24E-04 | 18.862 | 8.02E-05 |
| MGF360-6L | $p_2 = 0.016$, | $\omega_2 = 5.758$ | $p_1 = 0.027$, | $\omega = 4.210$ | 10.329 | 0.006 | 15.378 | 4.58E-04 |
| J5R /H108R | $p_2 = 0.169$, | $\omega_2 = 4.545$ | $p_1 = 0.169$, | $\omega = 4.545$ | 12.933 | 0.043 | 8.404 | 0.015 |
| A240L | $p_2 = 0.011$, | $\omega_2 = 7.581$ | $p_1 = 0.0176$, | $\omega = 5.558$ | 6.306 | 0.005 | 12.313 | 0.002 |
| MGF300-4L | $p_2 = 0.091$, | $\omega_2 = 2.927$ | $p_1 = 0.099$, | $\omega = 2.872$ | 11.274 | 0.004 | 11.088 | 0.004 |
| MGF360-12L | $p_2 = 0.015$, | $\omega_2 = 7.741$ | $p_1 = 0.015$, | $\omega = 7.723$ | 10.806 | 0.005 | 10.810 | 0.004 |
| P11.5/A137R | $p_2 = 0.177$, | $\omega_2 = 2.839$ | $p_1 = 0.177$, | $\omega = 2.843$ | 9.740 | 0.008 | 10.283 | 0.006 |
| MGF360-3L | $p_2 = 0.007$, | $\omega_2 = 9.342$ | $p_1 = 0.007$, | $\omega = 8.874$ | 7.301 | 0.026 | 8.288 | 0.016 |
| A238L | $p_2 = 0.005$, | $\omega_2 = 12.113$ | $p_1 = 0.005$, | $\omega = 11.777$ | 6.622 | 0.036 | 7.839 | 0.020 |
| B475L | $p_2 = 0.035$, | $\omega_2 = 3.895$ | $p_1 = 0.118$, | $\omega = 2.358$ | 7.786 | 0.020 | 7.750 | 0.021 |
| MGF505-5R | $p_2 = 0.035$, | $\omega_2 = 3.128$ | $p_1 = 0.046$, | $\omega = 2.905$ | 7.085 | 0.029 | 7.676 | 0.022 |
| P10/K78R | $p_2 = 0.060$, | $\omega_2 = 8.793$ | $p_1 = 0.060$, | $\omega = 8.856$ | 7.369 | 0.025 | 7.593 | 0.022 |
| L60L | $p_2 = 0.136$, | $\omega_2 = 2.763$ | $p_1 = 0.144$, | $\omega = 2.742$ | 4.954 | 0.084 | 7.567 | 0.023 |
| MGF360-8L | $p_2 = 0.217$, | $\omega_2 = 1.774$ | $p_1 = 0.216$, | $\omega = 1.779$ | 5.819 | 0.055 | 6.288 | 0.043 |
| MGF360-18R | $p_2 = 0.118$, | $\omega_2 = 2.123$ | $p_1 = 0.124$, | $\omega = 2.152$ | 5.416 | 0.067 | 6.947 | 0.031 |
| MGF505-1R | $p_2 = 0.015$, | $\omega_2 = 4.248$ | $p_1 = 0.023$, | $\omega = 3.667$ | 4.871 | 0.088 | 6.812 | 0.033 |
| MGF360-10L | $p_2 = 0.166$, | $\omega_2 = 1.352$ | $p_1 = 0.163$, | $\omega = 1.365$ | 1.517 | 0.468 | 6.942 | 0.031 |
| MGF360-4L | $p_2 = 0.041$, | $\omega_2 = 2.254$ | $p_1 = 0.083$, | $\omega = 1.921$ | 2.809 | 0.245 | 6.267 | 0.044 |
| Q706L | $p_2 = 0.002$, | $\omega_2 = 6.808$ | $p_1 = 0.002$, | $\omega = 6.861$ | 4.258 | 0.119 | 6.737 | 0.034 |

<sup>a</sup> LRT is the likelihood ratio test statistic calculated as $2\Delta l$ with $l$ the log likelihood for each model. The *p*-value was calculated using Chi-squared test.

**Table S7.** Three categories of proteins used for comparison of sequence variability.

| **Protein name** | **Antigenic?** | **Function** |
| --- | --- | --- |
| **Structural proteins not shown to be under positive selection** |  |  |
| P49/B438L | - | Minor capsid protein |
| P14.5/E120R | - | DNA-binding protein |
| P11.6/A104R | - | Histone-like DNA-binding protein |
| P22/KP177R | - | Inner envelop protein // virus entry |
| P12/O61R | - | Inner envelop protein |
| **Proteins shown to be antigenic in immunoassays** |  |  |
| MGF110-4L | + | Multigene family 110 |
| MGF110-5L | + | Multigene family 110 |
| C129R | + | Mn-dependent superoxide dismutase |
| E165R | + | dUTPase |
| E184L | + | Hypothetical protein |
| M448R | + | Microbody targeting signal-containing protein |
| F317L | + | Hypothetical protein |
| EP364R | + | ERCC4 nuclease domain |
| G1211R | + | DNA polymerase family B |
| NP1450L | + | RNA polymerase subunit 1 |
| E199L/J18L | + | Inner envelop protein // virus entry |
| P54/E183L | + | Inner envelop protein |
| P30/CP204L | + | Phosphoprotein binding to ribonucleoprotein K |
| P72/B464L | + | Major capsid protein |
| P17/D117L | + | Inner envelop protein |
| **Proteins previously shown to be involved in host-cell interactions** |  |  |
| A179L | - | Bcl 2 apoptosis inhibitor |
| I329L | + | Putative inhibitor of TLR3 signaling pathway |
| A224L | - | IAP apoptosis inhibitor |
| DP71L | + | Similar to herpes simplex virus ICP34.5 protein |
| L83L | - | Putative IL-1b binding protein |

**Table S8.**
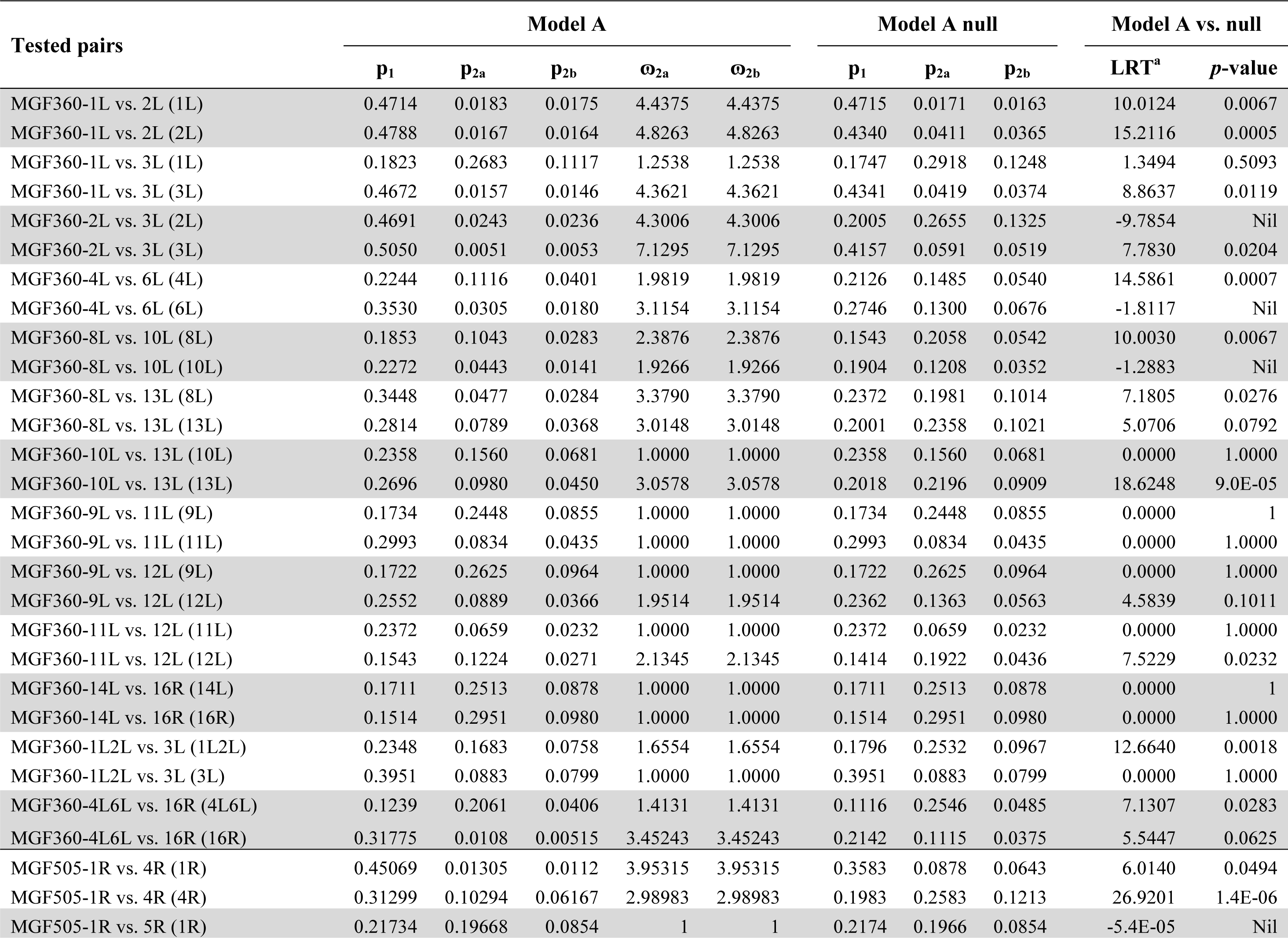
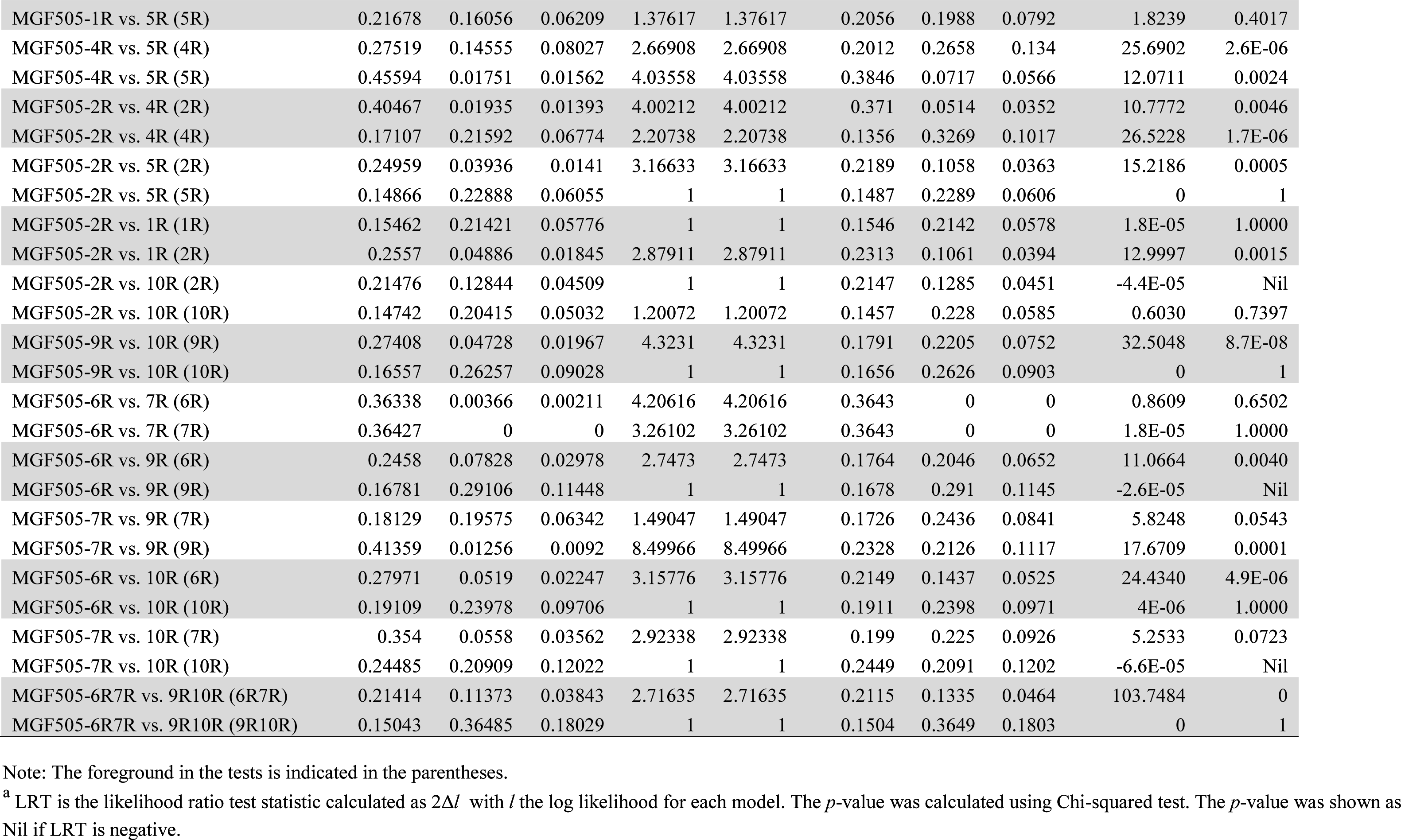
Pairs of paralogous genes/branches of MGF360 and MGF505 showing divergent selection at a fraction of sites based on the likelihood ratio tests of Model A of PAML.

| Tested pairs | Model A |  |  |  |  | Model A null |  |  | Model A vs. null |  |
| --- | --- | --- | --- | --- | --- | --- | --- | --- | --- | --- |
|  | P<sub>1</sub> | P<sub>2a</sub> | P<sub>2b</sub> | Ω<sub>2a</sub> | Ω<sub>2b</sub> | P<sub>1</sub> | P<sub>2a</sub> | P<sub>2b</sub> | LRT<sup>a</sup> | p-value |
| MGF360-1L vs. 2L (1L) | 0.4714 | 0.0183 | 0.0175 | 4.4375 | 4.4375 | 0.4715 | 0.0171 | 0.0163 | 10.0124 | 0.0067 |
| MGF360-1L vs. 2L (2L) | 0.4788 | 0.0167 | 0.0164 | 4.8263 | 4.8263 | 0.4340 | 0.0411 | 0.0365 | 15.2116 | 0.0005 |
| MGF360-1L vs. 3L (1L) | 0.1823 | 0.2683 | 0.1117 | 1.2538 | 1.2538 | 0.1747 | 0.2918 | 0.1248 | 1.3494 | 0.5093 |
| MGF360-1L vs. 3L (3L) | 0.4672 | 0.0157 | 0.0146 | 4.3621 | 4.3621 | 0.4341 | 0.0419 | 0.0374 | 8.8637 | 0.0119 |
| MGF360-2L vs. 3L (2L) | 0.4691 | 0.0243 | 0.0236 | 4.3006 | 4.3006 | 0.2005 | 0.2655 | 0.1325 | -9.7854 | Nil |
| MGF360-2L vs. 3L (3L) | 0.5050 | 0.0051 | 0.0053 | 7.1295 | 7.1295 | 0.4157 | 0.0591 | 0.0519 | 7.7830 | 0.0204 |
| MGF360-4L vs. 6L (4L) | 0.2244 | 0.1116 | 0.0401 | 1.9819 | 1.9819 | 0.2126 | 0.1485 | 0.0540 | 14.5861 | 0.0007 |
| MGF360-4L vs. 6L (6L) | 0.3530 | 0.0305 | 0.0180 | 3.1154 | 3.1154 | 0.2746 | 0.1300 | 0.0676 | -1.8117 | Nil |
| MGF360-8L vs. 10L (8L) | 0.1853 | 0.1043 | 0.0283 | 2.3876 | 2.3876 | 0.1543 | 0.2058 | 0.0542 | 10.0030 | 0.0067 |
| MGF360-8L vs. 10L (10L) | 0.2272 | 0.0443 | 0.0141 | 1.9266 | 1.9266 | 0.1904 | 0.1208 | 0.0352 | -1.2883 | Nil |
| MGF360-8L vs. 13L (8L) | 0.3448 | 0.0477 | 0.0284 | 3.3790 | 3.3790 | 0.2372 | 0.1981 | 0.1014 | 7.1805 | 0.0276 |
| MGF360-8L vs. 13L (13L) | 0.2814 | 0.0789 | 0.0368 | 3.0148 | 3.0148 | 0.2001 | 0.2358 | 0.1021 | 5.0706 | 0.0792 |
| MGF360-10L vs. 13L (10L) | 0.2358 | 0.1560 | 0.0681 | 1.0000 | 1.0000 | 0.2358 | 0.1560 | 0.0681 | 0.0000 | 1.0000 |
| MGF360-10L vs. 13L (13L) | 0.2696 | 0.0980 | 0.0450 | 3.0578 | 3.0578 | 0.2018 | 0.2196 | 0.0909 | 18.6248 | 9.0E-05 |
| MGF360-9L vs. 11L (9L) | 0.1734 | 0.2448 | 0.0855 | 1.0000 | 1.0000 | 0.1734 | 0.2448 | 0.0855 | 0.0000 | 1 |
| MGF360-9L vs. 11L (11L) | 0.2993 | 0.0834 | 0.0435 | 1.0000 | 1.0000 | 0.2993 | 0.0834 | 0.0435 | 0.0000 | 1.0000 |
| MGF360-9L vs. 12L (9L) | 0.1722 | 0.2625 | 0.0964 | 1.0000 | 1.0000 | 0.1722 | 0.2625 | 0.0964 | 0.0000 | 1.0000 |
| MGF360-9L vs. 12L (12L) | 0.2552 | 0.0889 | 0.0366 | 1.9514 | 1.9514 | 0.2362 | 0.1363 | 0.0563 | 4.5839 | 0.1011 |
| MGF360-11L vs. 12L (11L) | 0.2372 | 0.0659 | 0.0232 | 1.0000 | 1.0000 | 0.2372 | 0.0659 | 0.0232 | 0.0000 | 1.0000 |
| MGF360-11L vs. 12L (12L) | 0.1543 | 0.1224 | 0.0271 | 2.1345 | 2.1345 | 0.1414 | 0.1922 | 0.0436 | 7.5229 | 0.0232 |
| MGF360-14L vs. 16R (14L) | 0.1711 | 0.2513 | 0.0878 | 1.0000 | 1.0000 | 0.1711 | 0.2513 | 0.0878 | 0.0000 | 1 |
| MGF360-14L vs. 16R (16R) | 0.1514 | 0.2951 | 0.0980 | 1.0000 | 1.0000 | 0.1514 | 0.2951 | 0.0980 | 0.0000 | 1.0000 |
| MGF360-1L2L vs. 3L (1L2L) | 0.2348 | 0.1683 | 0.0758 | 1.6554 | 1.6554 | 0.1796 | 0.2532 | 0.0967 | 12.6640 | 0.0018 |
| MGF360-1L2L vs. 3L (3L) | 0.3951 | 0.0883 | 0.0799 | 1.0000 | 1.0000 | 0.3951 | 0.0883 | 0.0799 | 0.0000 | 1.0000 |
| MGF360-4L6L vs. 16R (4L6L) | 0.1239 | 0.2061 | 0.0406 | 1.4131 | 1.4131 | 0.1116 | 0.2546 | 0.0485 | 7.1307 | 0.0283 |
| MGF360-4L6L vs. 16R (16R) | 0.31775 | 0.0108 | 0.00515 | 3.45243 | 3.45243 | 0.2142 | 0.1115 | 0.0375 | 5.5447 | 0.0625 |
| MGF505-1R vs. 4R (1R) | 0.45069 | 0.01305 | 0.0112 | 3.95315 | 3.95315 | 0.3583 | 0.0878 | 0.0643 | 6.0140 | 0.0494 |
| MGF505-1R vs. 4R (4R) | 0.31299 | 0.10294 | 0.06167 | 2.98983 | 2.98983 | 0.1983 | 0.2583 | 0.1213 | 26.9201 | 1.4E-06 |
| MGF505-1R vs. 5R (1R) | 0.21734 | 0.19668 | 0.0854 | 1 | 1 | 0.2174 | 0.1966 | 0.0854 | -5.4E-05 | Nil |
| MGF505-1R vs. 5R (5R) | 0.21678 | 0.16056 | 0.06209 | 1.37617 | 1.37617 | 0.2056 | 0.1988 | 0.0792 | 1.8239 | 0.4017 |
| MGF505-4R vs. 5R (4R) | 0.27519 | 0.14555 | 0.08027 | 2.66908 | 2.66908 | 0.2012 | 0.2658 | 0.134 | 25.6902 | 2.6E-06 |
| MGF505-4R vs. 5R (5R) | 0.45594 | 0.01751 | 0.01562 | 4.03558 | 4.03558 | 0.3846 | 0.0717 | 0.0566 | 12.0711 | 0.0024 |
| MGF505-2R vs. 4R (2R) | 0.40467 | 0.01935 | 0.01393 | 4.00212 | 4.00212 | 0.371 | 0.0514 | 0.0352 | 10.7772 | 0.0046 |
| MGF505-2R vs. 4R (4R) | 0.17107 | 0.21592 | 0.06774 | 2.20738 | 2.20738 | 0.1356 | 0.3269 | 0.1017 | 26.5228 | 1.7E-06 |
| MGF505-2R vs. 5R (2R) | 0.24959 | 0.03936 | 0.0141 | 3.16633 | 3.16633 | 0.2189 | 0.1058 | 0.0363 | 15.2186 | 0.0005 |
| MGF505-2R vs. 5R (5R) | 0.14866 | 0.22888 | 0.06055 | 1 | 1 | 0.1487 | 0.2289 | 0.0606 | 0 | 1 |
| MGF505-2R vs. 1R (1R) | 0.15462 | 0.21421 | 0.05776 | 1 | 1 | 0.1546 | 0.2142 | 0.0578 | 1.8E-05 | 1.0000 |
| MGF505-2R vs. 1R (2R) | 0.2557 | 0.04886 | 0.01845 | 2.87911 | 2.87911 | 0.2313 | 0.1061 | 0.0394 | 12.9997 | 0.0015 |
| MGF505-2R vs. 10R (2R) | 0.21476 | 0.12844 | 0.04509 | 1 | 1 | 0.2147 | 0.1285 | 0.0451 | -4.4E-05 | Nil |
| MGF505-2R vs. 10R (10R) | 0.14742 | 0.20415 | 0.05032 | 1.20072 | 1.20072 | 0.1457 | 0.228 | 0.0585 | 0.6030 | 0.7397 |
| MGF505-9R vs. 10R (9R) | 0.27408 | 0.04728 | 0.01967 | 4.3231 | 4.3231 | 0.1791 | 0.2205 | 0.0752 | 32.5048 | 8.7E-08 |
| MGF505-9R vs. 10R (10R) | 0.16557 | 0.26257 | 0.09028 | 1 | 1 | 0.1656 | 0.2626 | 0.0903 | 0 | 1 |
| MGF505-6R vs. 7R (6R) | 0.36338 | 0.00366 | 0.00211 | 4.20616 | 4.20616 | 0.3643 | 0 | 0 | 0.8609 | 0.6502 |
| MGF505-6R vs. 7R (7R) | 0.36427 | 0 | 0 | 3.26102 | 3.26102 | 0.3643 | 0 | 0 | 1.8E-05 | 1.0000 |
| MGF505-6R vs. 9R (6R) | 0.2458 | 0.07828 | 0.02978 | 2.7473 | 2.7473 | 0.1764 | 0.2046 | 0.0652 | 11.0664 | 0.0040 |
| MGF505-6R vs. 9R (9R) | 0.16781 | 0.29106 | 0.11448 | 1 | 1 | 0.1678 | 0.291 | 0.1145 | -2.6E-05 | Nil |
| MGF505-7R vs. 9R (7R) | 0.18129 | 0.19575 | 0.06342 | 1.49047 | 1.49047 | 0.1726 | 0.2436 | 0.0841 | 5.8248 | 0.0543 |
| MGF505-7R vs. 9R (9R) | 0.41359 | 0.01256 | 0.0092 | 8.49966 | 8.49966 | 0.2328 | 0.2126 | 0.1117 | 17.6709 | 0.0001 |
| MGF505-6R vs. 10R (6R) | 0.27971 | 0.0519 | 0.02247 | 3.15776 | 3.15776 | 0.2149 | 0.1437 | 0.0525 | 24.4340 | 4.9E-06 |
| MGF505-6R vs. 10R (10R) | 0.19109 | 0.23978 | 0.09706 | 1 | 1 | 0.1911 | 0.2398 | 0.0971 | 4E-06 | 1.0000 |
| MGF505-7R vs. 10R (7R) | 0.354 | 0.0558 | 0.03562 | 2.92338 | 2.92338 | 0.199 | 0.225 | 0.0926 | 5.2533 | 0.0723 |
| MGF505-7R vs. 10R (10R) | 0.24485 | 0.20909 | 0.12022 | 1 | 1 | 0.2449 | 0.2091 | 0.1202 | -6.6E-05 | Nil |
| MGF505-6R7R vs. 9R10R (6R7R) | 0.21414 | 0.11373 | 0.03843 | 2.71635 | 2.71635 | 0.2115 | 0.1335 | 0.0464 | 103.7484 | 0 |
| MGF505-6R7R vs. 9R10R (9R10R) | 0.15043 | 0.36485 | 0.18029 | 1 | 1 | 0.1504 | 0.3649 | 0.1803 | 0 | 1 |

Note: The foreground in the tests is indicated in the parentheses.
<sup>a</sup> LRT is the likelihood ratio test statistic calculated as $2\Delta l$ with $l$ the log likelihood for each model. The $p$ -value was calculated using Chi-squared test. The $p$ -value was shown as Nil if LRT is negative.

**Fig. S1.**
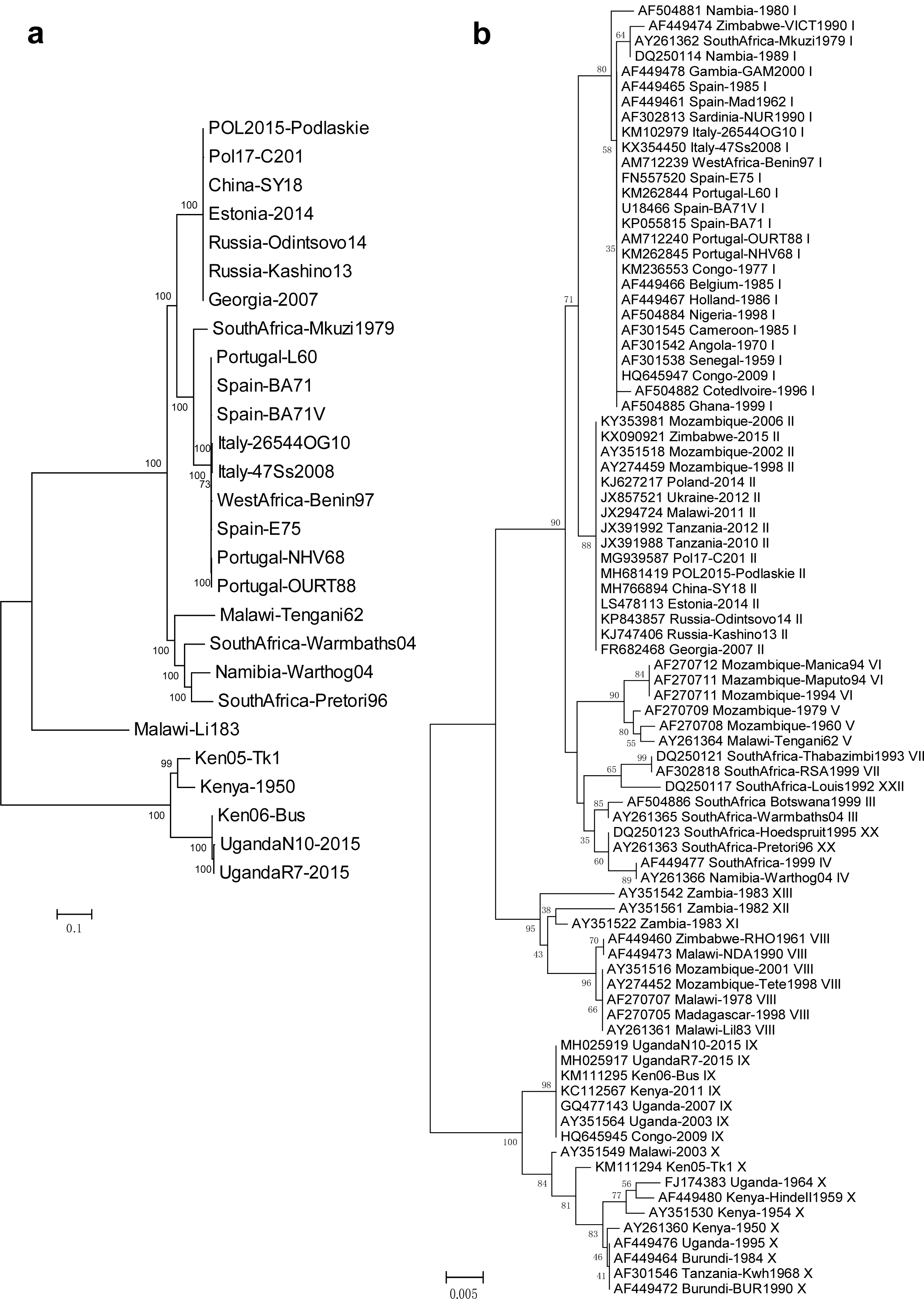
Phylogenetic structure constructed for the core genome of ASFV (a) and the C-terminal 414 bp of the structural gene p72 presented in a dendrogram tree (b). The tree for the p72 was built from the isolates compiled from the NCBI database https://www.ncbi.nlm.nih.gov/. A total of 85 non-redundant isolates were obtained with unique geographical location and isolate time and were used for tree construction. The tree was inferred using the Neighboring-Joining method with 1000 bootstrap without consensus. The isolate names in the p72 tree were presented as the combination of accession number, location, time and genotype.

**Fig. S2.**
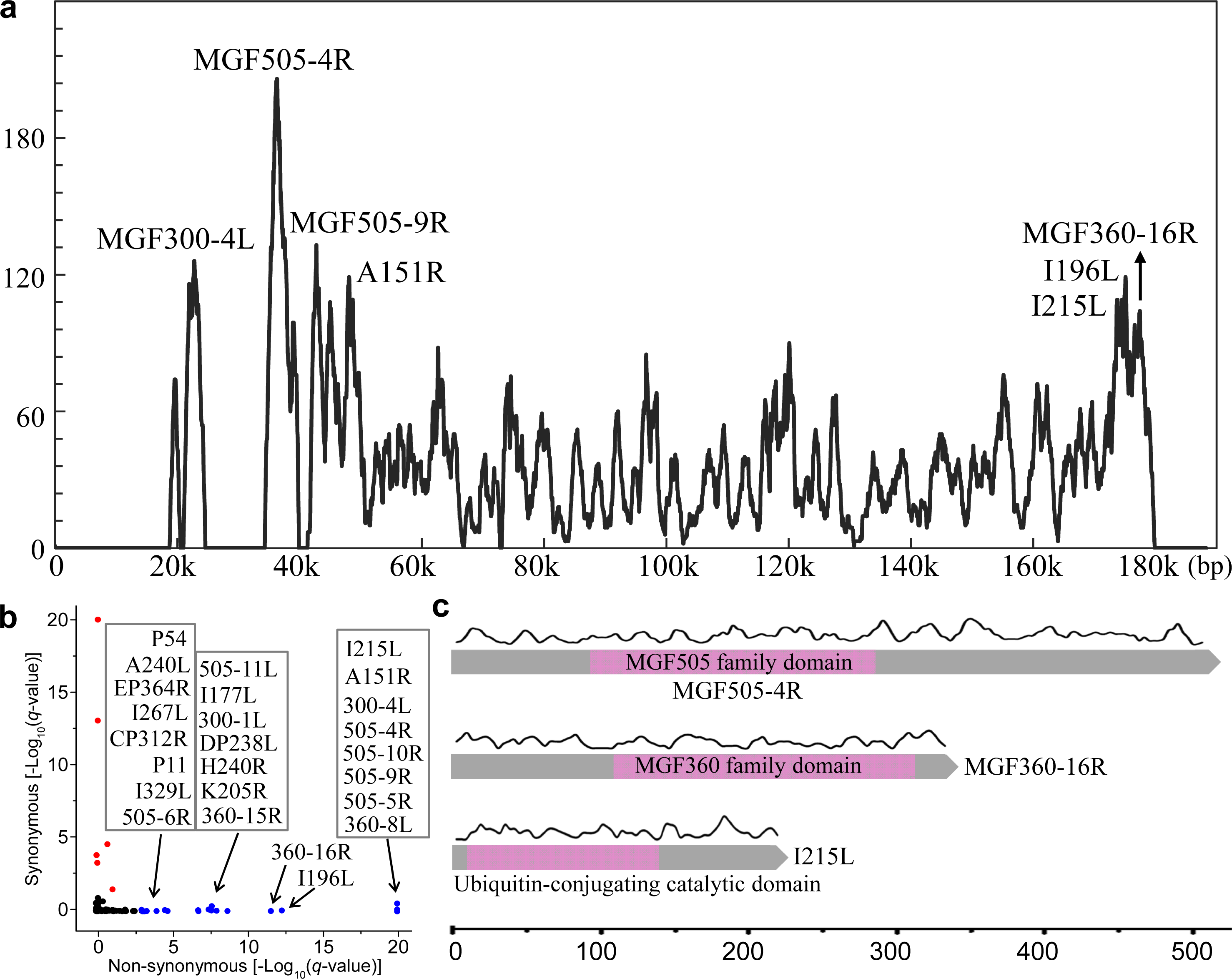
Profiling of the distribution of non-synonymous mutations along the ASFV genomes. (a) The density distribution (number of mutations per kb) of non-synonymous mutations along the genome of the representative strain Georgia-2007. The top genes with the highest density of non-synonymous mutations are indicated. (b) All genes enriched with non-synonymous mutations (*q*-value ≤ 0.001) but not with synonymous mutations (*q*-value > 0.05) are shown in blue dots. The genes enriched with synonymous mutations (*q*-value ≤ 0.05) but not with non-synonymous mutations (*q*-value > 0.05) are shown in red dots. The genes are not enriched with either mutations are in black dots. The *q*-value is defined as the multiple testing corrected *p*-value using the Benjamini-Hochberg procedure. The *p*-value was calculated with the Hypergeometric test. (c) A detailed view of the density distribution of non-synonymous mutations for three top genes is depicted along the domain architecture of the genes. There is no significant difference of the mutation distribution between different functional domains.

**Fig. S3.**
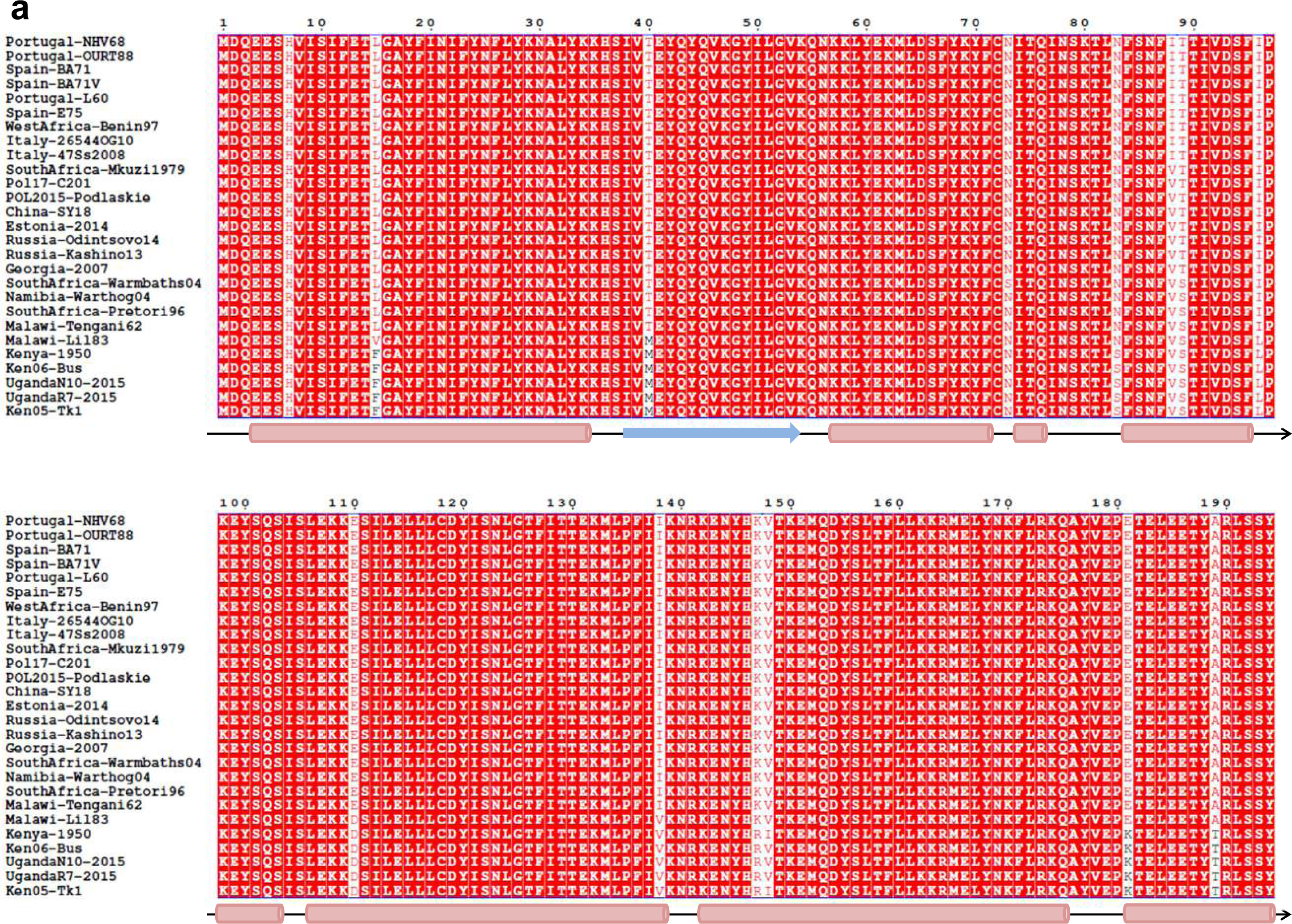
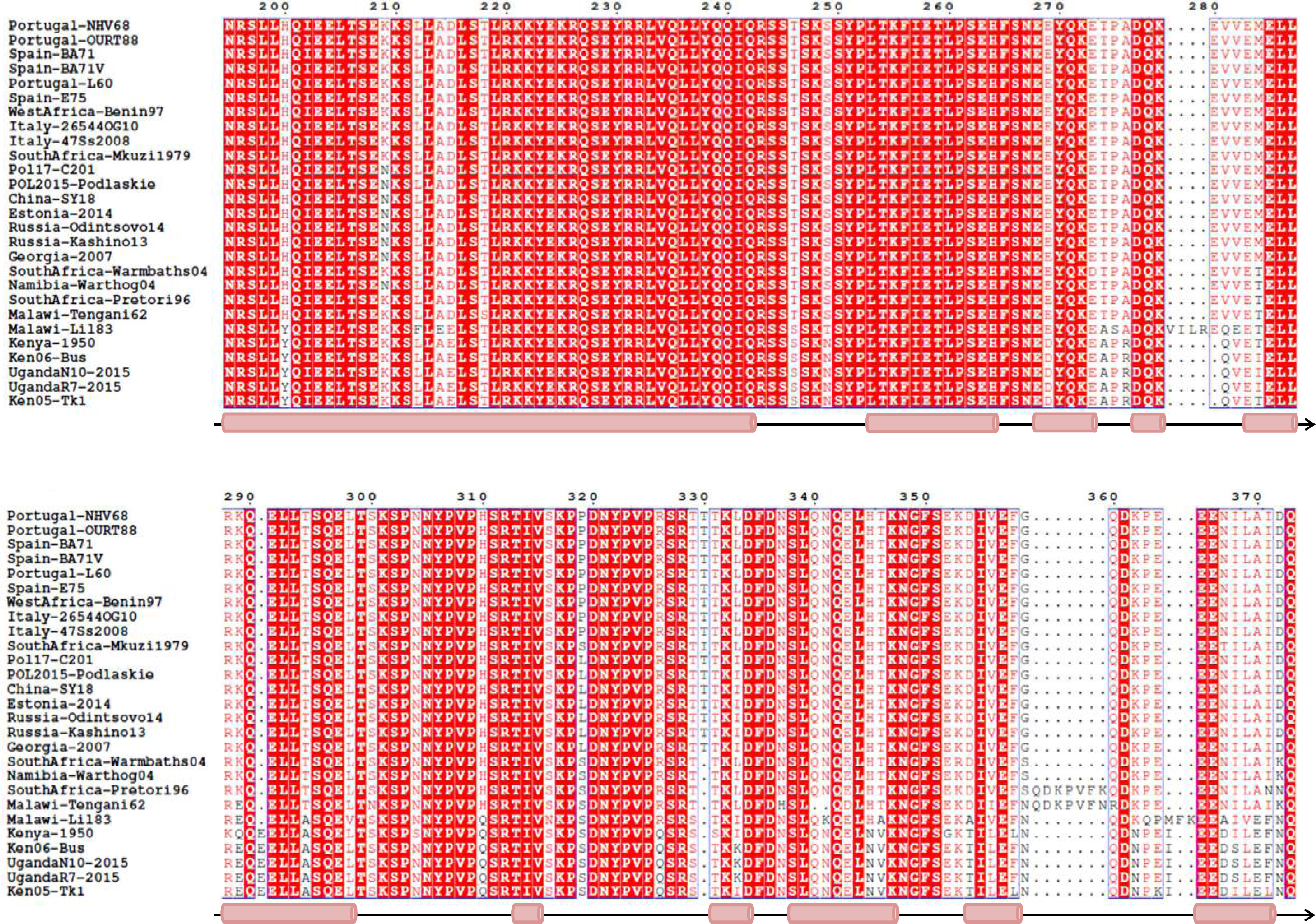
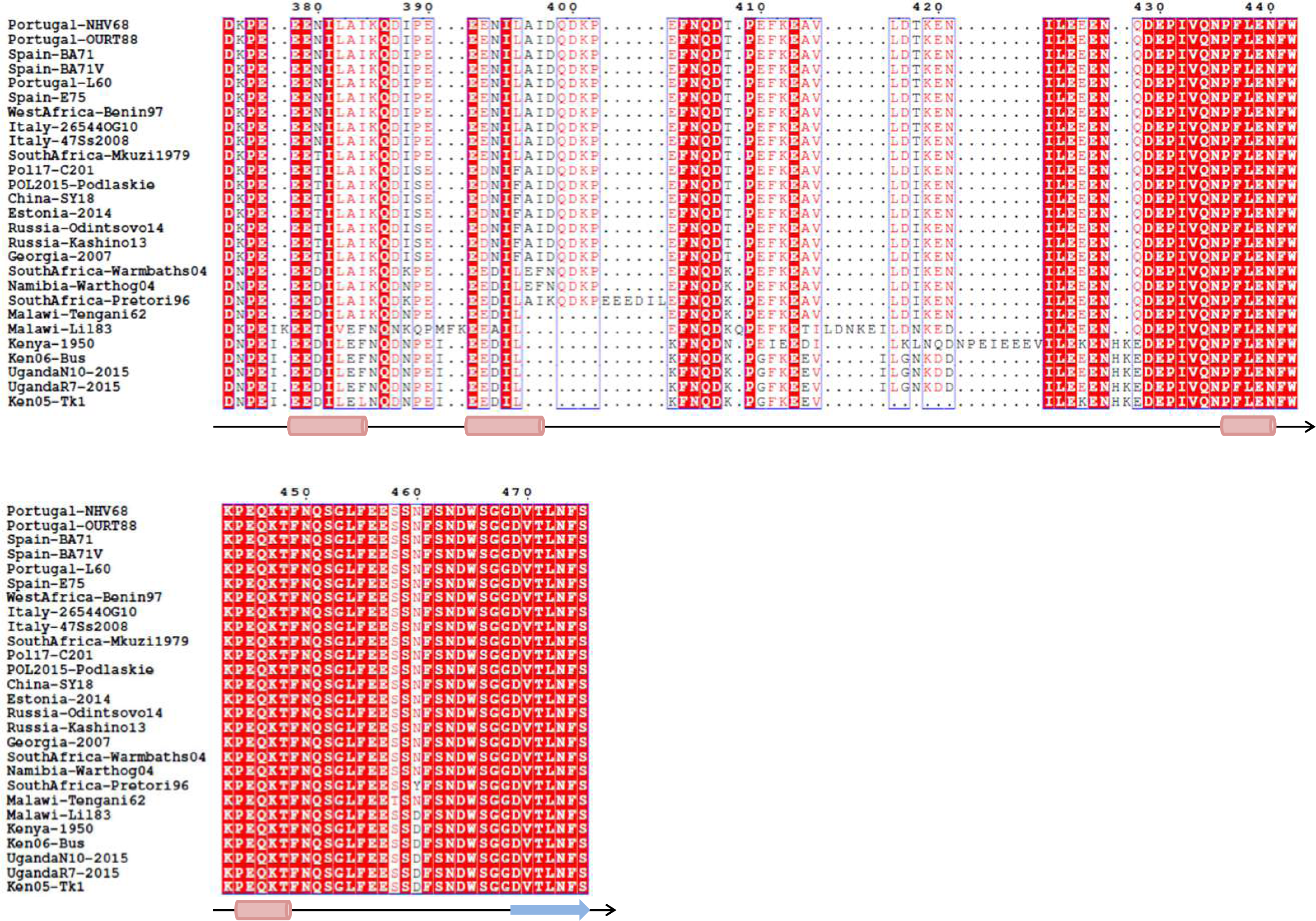
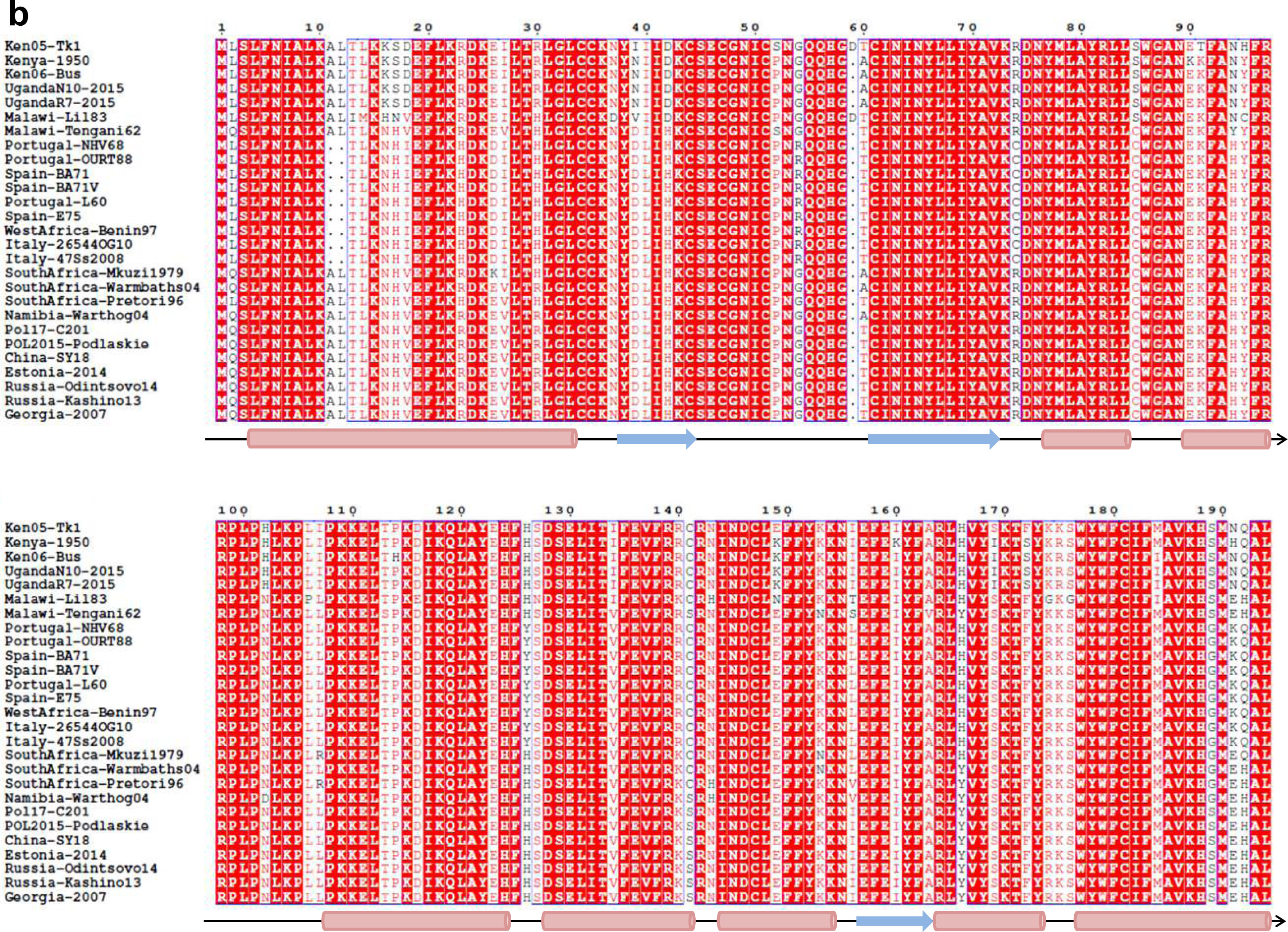
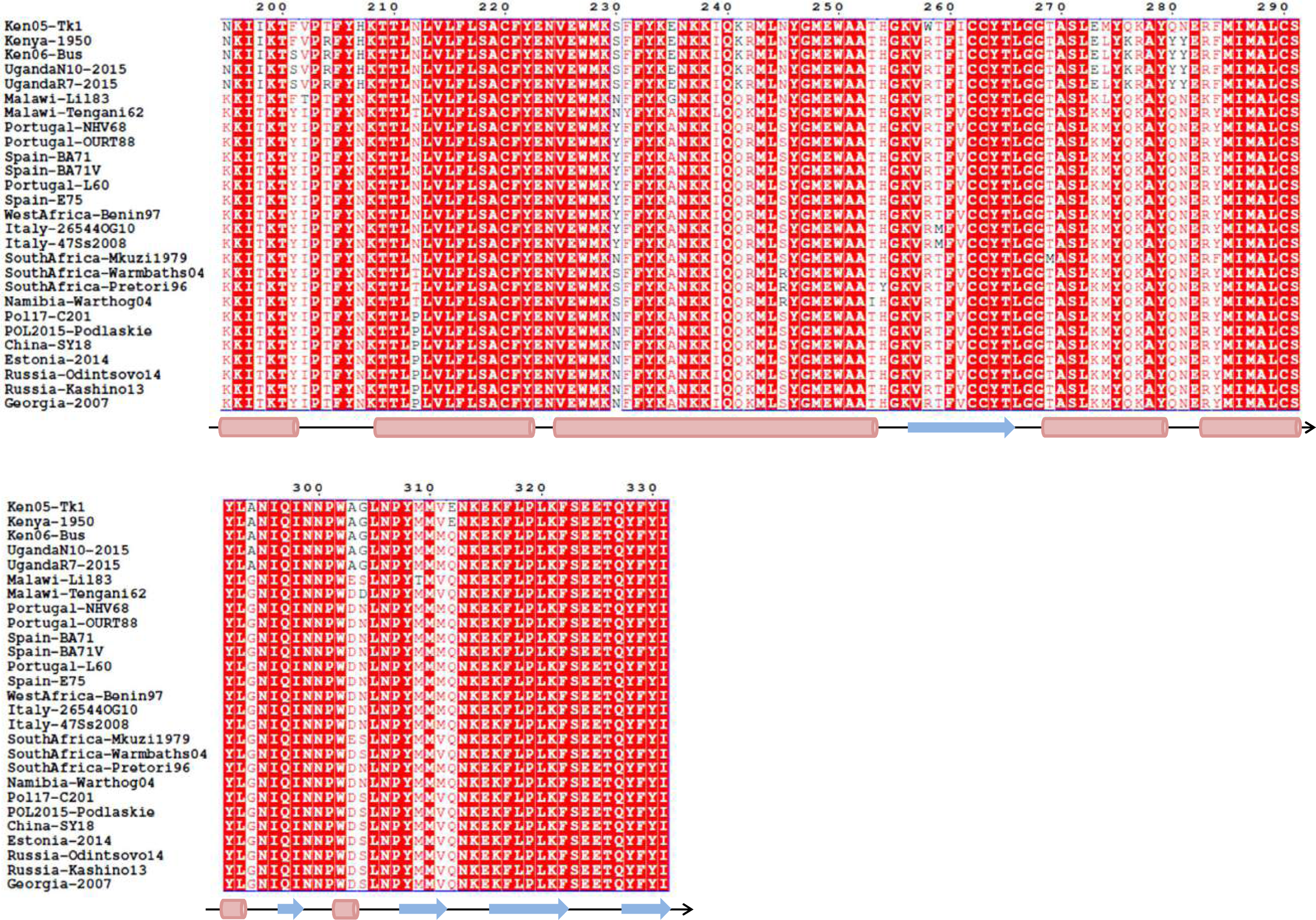
The predicted secondary structures of B475L (a) and MGF300-4L (b. The secondary structures are represented as α-helices (cylinders), β-strands (arrows), or coiled loops (lines). Both proteins are predominated by tandem α-helices.

**Fig. S4.**
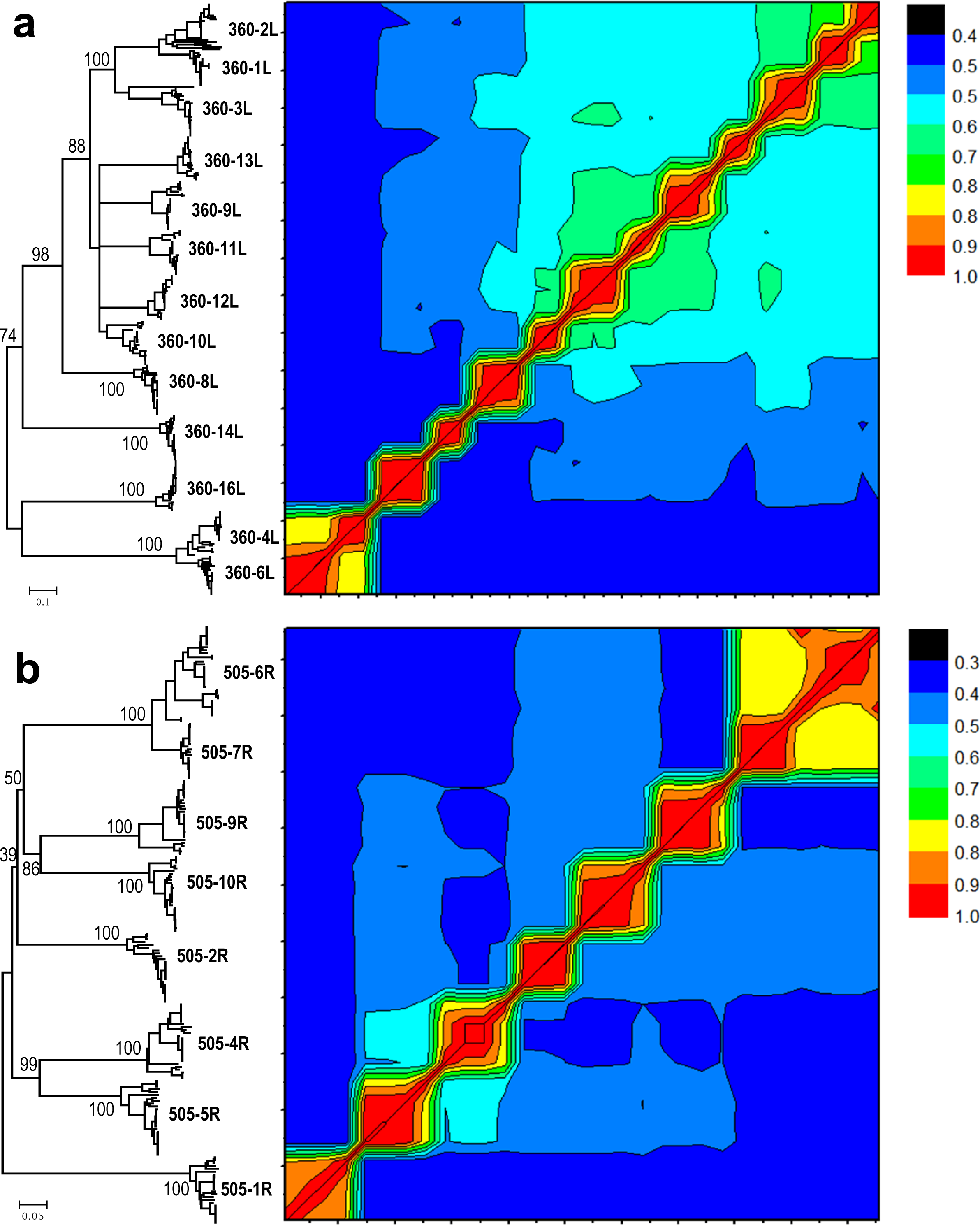
The heatmap of pair-wise nucleotide similarities of the orthologous and paralogous genes of MGF360 (a) and MGF505 (b) along with the phylogenetic structure. The phylogenetic structure was inferred using Neighbor-Joining method with 1000 bootstraps for the orthologs and paralogs for each member of MGF360 and MGF505. The sequence alignments used for the phylogeny are provided in the public link https://figshare.com/projects/ASFV_alignment/82718 (see “Data availability”). Only nodes with the support value > 30 in the phylogeny are shown. A colored scale for the nucleotide similarities is given on the right side of the heatmap. The similarities between orthologous genes are much higher than that for paralogous members, and therefore the former cluster together in the trees, except three isolates of MGF360-1L (from Kenya-1950, Ken05-Tk1, and Spain-E75), which cluster together with MGF360-2L, and five isolate of MGF505-7R (from Malawi-Lil83, Kenya-1950, Ken05-Tk1, Ken06-Bus, and UgandaN10-2015), which cluster together with MGF505-6R.

**Fig. S5.**
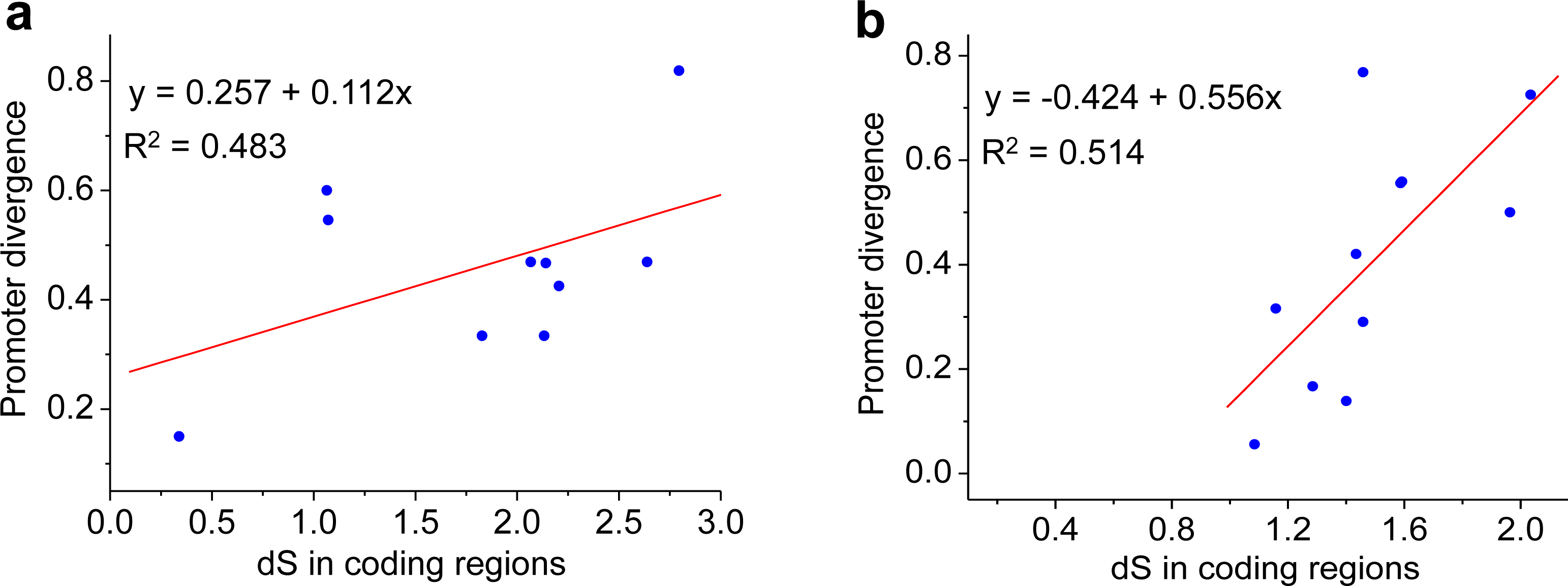
Correlation between the promoter divergence (y axis) and the synonymous substitution rate for each pair of genes/branches (x axis) in MGF360 (a) and MGF505 (b). The fitted lines of linear regression are shown in red and the fitting equation and Pearson correlations R^2^ are indicated.

